# Brain composition in *Heliconius* butterflies, post-eclosion growth and experience dependent neuropil plasticity

**DOI:** 10.1101/017913

**Authors:** Stephen H. Montgomery, Richard M. Merrill, Swidbert R. Ott

## Abstract

Behavioral and sensory adaptations are often based in the differential expansion of brain components. These volumetric differences represent changes in investment, processing capacity and/or connectivity, and can be used to investigate functional and evolutionary relationships between different brain regions, and between brain composition and behavioral ecology. Here, we describe the brain composition of two species of *Heliconius* butterflies, a long-standing study system for investigating ecological adaptation and speciation. We confirm a previous report of striking mushroom body expansion, and explore patterns of post-eclosion growth and experience-dependent plasticity in neural development. This analysis uncovers age-and experience-dependent post-emergence mushroom body growth comparable to that in foraging hymenoptera, but also identifies plasticity in several other neuropil. An interspecific analysis indicates that *Heliconius* display remarkable levels of investment in mushroom bodies for a lepidopteran, and indeed rank highly compared to other insects. Our analyses lay the foundation for future comparative and experimental analyses that will establish *Heliconius* as a useful case study in evolutionary neurobiology.

## INTRODUCTION

Behavioral adaptations are largely based in changes in brain function. In some cases this includes differential expansion of individual brain structures, or functionally related systems, that betray underlying changes in neuron number or circuitry. These provide an opportunity to study the neural basis of adaptive behavior, particularly in clades with known ecological specializations. The Neotropical genus *Heliconius* (Heliconiinae, Nymphalidae) display a number of striking behavioral adaptations including a dietary adaptation unique among Lepidoptera; adult pollen feeding (Gilbert, 1972, 1975). With the exception of four species formerly ascribed to the genus *Neruda* (Beltrán et al., 2007; Kozak et al., 2015), all *Heliconius* actively collect and ingest pollen as adults. This provides a source of amino acids and permits a greatly extended lifespan of up to six months without reproductive senescence (Gilbert, 1972; Benson, 1972; Ehrlich and Gilbert, 1973). Without access to pollen *Heliconius* suffer a major reduction in longevity and reproductive success (Gilbert, 1972; Dunlap-Pianka et al., 1977; O’Brien et al., 2003).

Several lines of evidence suggest selection for pollen feeding has shaped *Heliconius* foraging behavior. Pollen is collected from a restricted range of mostly Cucurbitaceous plants (Estrada and Jiggins, 2002), which occur at low densities (Gilbert, 1975). Individuals inhabit home ranges of typically less than 1 km^2^, within which they repeatedly utilize a small number of roosting sites that they return to with high fidelity (Turner, 1971; Benson, 1972; Gilbert, 1975; Mallet, 1986; Murawski and Gilbert, 1986; Finkbeiner, 2014). On leaving the roost individuals visit feeding sites with a level of consistency in time and space that strongly suggests ‘trap-lining’ behavior (Ehrlich and Gilbert, 1973; Gilbert, 1975, 1993; Mallet, 1986), analogous to that observed in foraging bees (Janzen, 1971; Heinrich, 1979). Roosts themselves are located visually (Jones, 1930; Gilbert, 1972; Ehrlich and Gilbert, 1973; Mallet, 1986), and older individuals tend to be more efficient foragers (Boggs et al., 1981; Gilbert, 1993). Together these observations suggest the evolution of pollen feeding in *Heliconius* was facilitated by an enhanced, visually-orientated and time-compensated memory that utilizes long distance landmarks (Gilbert, 1975). The evolution of this behavior must involve “some elaboration of the nervous system” (Turner, 1981). This elaboration is suggested to occur in the mushroom bodies, which Sivinski (1989) reported are 3–4 times larger in *Heliconius charithonia* than in six other species of butterfly, including two non-pollen feeding Heliconiini.

Insect mushroom bodies have a variety of roles in olfactory associative learning, sensory integration, filtering and attention (Zars, 2000; Farris, 2005, 2013; Menzel, 2014). Direct experimental evidence suggests the mushroom bodies mediate place memory in *Periplaneta americana* (Mizunami et al., 1998; Lent et al., 2007). Two lines of indirect evidence further implicate the mushroom bodies in allocentric memory in other insects. First, comparisons across species suggest that extreme evolutionary expansion of the mushroom body may commonly be associated with changes in foraging behavior that depend on spatial memory or the complexity of sensory information utilized by the species (Farris, 2005, 2013). For example, phylogenetic comparisons across Hymenoptera demonstrate the volumetric expansion and elaboration of the Euhymenopteran mushroom body occurs coincidently with the origin of parasitoidism (Farris and Schulmeister, 2011), a behavioral adaptation that involves place-centered foraging and spatial memory for host location (Rosenheim, 1987; van Nouhuys and Kaartinen, 2008). Second, mushroom bodies are ontogenetically plastic, and this plasticity has been linked to foraging behavior. Again, the trap-lining Hymenoptera illustrate the link between foraging behavior and the mushroom bodies (Withers et al., 1993; Durst et al., 1994; Capaldi et al., 1999; Farris et al., 2001). Honeybees show two forms of post-eclosion growth in mushroom body volume; age dependent growth, which occurs regardless of environmental variation, and experience dependent growth which increases with foraging or social experience (Withers et al., 1993; Durst et al., 1994; Fahrbach et al., 1998, 2003; Farris et al., 2001; Maleszka et al., 2009). In other Hymenoptera there is close correspondence between the rate and timing of mushroom body growth and the onset of foraging behavior (Gronenberg et al., 1996; Kühn-Bühlmann and Wehner, 2006; Withers et al., 2008; Jones et al., 2013). Whether the trap-lining behavior observed in *Heliconius* is associated with similar ontogenetic plasticity is not known.

Here we confirm Sivinksi’s (1989) observation of a phylogenetic expansion of the mushroom bodies in *Heliconius*. We further demonstrate ontogenetic and environmentally induced plasticity comparable in size to trap-lining Hymenoptera. Together these analyses suggest the mushroom bodies may have a role in the allocentric spatial foraging observed in *Heliconius*, and lay the groundwork for comparative analyses across Heliconiini examining the origin and timing of mushroom body expansion.

## MATERIALS & METHODS

### Animals

We collected five males and five females of two species of *Heliconius*, *H. hecale melicerta* and *H. erato demophoon* from wild populations around Gamboa (9°7.4′ N, 79°42.2′ W, elevation 60 m) and the nearby Soberanía National Park, República de Panamá. We assume all wild-caught individuals were sexually mature, and that the age range is not biased between species or sexes. Wild individuals were compared to individuals from first or second-generation insectary-reared stock populations, descended from wild caught parents from the same sampling localities. Stock populations were kept in controlled conditions in cages (c. 1 × 2 × 2 m) of mixed sex at roughly equal densities. Cages were housed at the *Heliconius* insectaries at the Smithsonian Tropical Research Institute’s (STRI) facility in Gamboa. Stocks had access to their preferred host plant (*Passiflora biflora* and *P. vitifolia* respectively for *H. erato* and *H. hecale*), a pollen source (*Psychotria elata*) and feeders containing c. 20% sugar solution with an additional bee-pollen supplement to ensure an excess of pollen. Larvae were allowed to feed naturally on the host plant.

After emergence from the pupae insectary-reared individuals were collected for two age groups, a recently emerged ‘young’ group (1–3 days post emergence) and an ‘old’ group (2–3 weeks post emergence). *Heliconius* undergo a “callow” period of general inactivity immediately after emergence that lasts about 5 days, during which flight behavior is weak and males are sexually inactive (Mallet, 1980). These age groups therefore represent behaviorally immature and mature individuals. For *H. hecale* 5 males and 5 females were sampled for both age groups, in *H. erato* 4 males and 6 females were sampled for the ‘young’ group and 5 males and 4 females were sampled for the ‘old’ group. In samples for which the exact time of emergence was known there was no significant difference between *H. hecale* and *H. erato* in age structure of the old (*H. erato*: mean = 22.6 days, SD = 8.6; *H. hecale*: mean = 26.4 days, SD = 5.5; *t*_*13*_= -0.899, *p* = 0.385) or young (*H. erato*: mean = 1.7 days, SD = 0.8; *H. hecale*: mean = 1.3 days, SD = 1.1; *t*_17_= 0.829, *p* = 0.419) insectary-reared groups. Three body size measurements were taken for each individual: body mass, weighted to 0.01 g using a OHAUS pocket balance (model YA102), body length, and wingspan, measured using FreeLOGIX digital calipers. Samples were collected and exported under permits SEX/A-3-12 and SE/A-7-13 obtained from the Autoridad Nacional del Ambiente, República de Panamá in conjunction with STRI.

### Antibodies and sera for neuropil staining

We used indirect immunofluorescence staining against synapsin to reveal the neuropil structure of the brain under a confocal microscope (Ott, 2008). This technique exploits the abundant expression of synapsin, a vesicle-associated protein, at presynaptic sites. Monoclonal mouse anti-synapsin antibody 3C11 (anti-SYNORF1; (Klagges et al., 1996) was obtained from the Developmental Studies Hybridoma Bank (DSHB), University of Iowa, Department of Biological Sciences, Iowa City, IA 52242, USA (RRID: AB_2315424). The 3C11 antibody was raised against a bacterially expressed fusion protein generated by adding a glutathione S-transferase (GST)-tag to cDNA comprised of most of the 5′ open reading frame 1 of the *Drosophila melanogaster* synapsin gene (*Syn,* CG3985). The binding specificity of this antibody was characterised in *D. melanogaster* (Klagges et al., 1996) and confirmed in synapsin null mutants by Godenschwege et al. (2004). The epitope was later narrowed down to within LFGGMEVCGL in the C domain (Hofbauer et al., 2009). Bioinformatic analysis has confirmed the presence of this motif in lepidopteran genomes, and demonstrated that it is highly conserved across Lepidoptera (Montgomery and Ott, 2015). Binding specificity in *M. sexta* has been confirmed by western blot analysis (Utz et al., 2008) and 3C11 immunostaining has been used as an anatomical marker of synaptic neuropil in a wide range of arthropod species including several Lepidoptera: *D. plexippus* (Heinze and Reppert, 2012), *G. zavaleta* (Montgomery and Ott, 2015), *H. virescens* (Kvello et al., 2009) and *M. sexta* (El Jundi et al., 2009). Cy2-conjugated affinity-purified polyclonal goat anti-mouse IgG (H+L) antibody (Jackson ImmunoResearch Laboratories, West Grove, PA) was obtained from Stratech Scientific Ltd., Newmarket, Suffolk, UK (Jackson ImmunoResearch Cat No. 115-225-146, RRID: AB_2307343).

### Immunocytochemistry

Brains were fixed and stained following a published protocol (Ott, 2008). The protocol was divided into two stages, the first of which was performed at the STRI Gamboa Field Station. The brain was exposed under HEPES-buffered saline (HBS; 150 mM NaCl; 5 mM KCl; 5 mM CaCl_2_; 25 mM sucrose; 10 mM HEPES; pH 7.4) and fixed in situ for 16–20 hours at room temperature (RT) in zinc-formaldehyde solution (ZnFA; 0.25% (18.4 mM) ZnCl_2_; 0.788% (135 mM) NaCl; 1.2% (35 mM) sucrose; 1% formaldehyde) under agitation. The brain was subsequently dissected out under HBS, washed (3 × in HBS), placed into 80% methanol/20% DMSO for 2 hours under agitation, transferred to 100% methanol and stored at RT. After transportation to the UK samples were stored at -20°C.

In the second stage of the protocol the samples were brought to RT and rehydrated in a decreasing methanol series (90%, 70%, 50%, 30%, 0% in 0.1 M Tris buffer, pH 7.4, 10 minutes each). Normal goat serum (NGS; New England BioLabs, Hitchin, Hertfordshire, UK) and antibodies were diluted in 0.1 M phosphate-buffered saline (PBS; pH 7.4) containing 1% DMSO and 0.005% NaN_3_ (PBSd). After a pre-incubation in 5% NGS (PBSd-NGS) for 2 hours at RT, antibody 3C11 was applied at a 1:30 dilution in PBSd-NGS for 3.5 days at 4°C under agitation. The brains were rinsed in PBSd (3 × 2 hours) before applying the Cy2-conjugated anti-mouse antibody 1:100 in PBSd-NGS for 2.5 days at 4°C under agitation. This was followed by increasing concentrations of glycerol (1%, 2%, 4% for 2 hours each, 8%, 15%, 30%, 50%, 60%, 70% and 80% for 1 hour each) in 0.1 M Tris buffer with DMSO to 1%. The brains were then passed in a drop of 80% glycerol directly into 100% ethanol. After agitation for 30 minutes the ethanol was refreshed (3 × 30 minute incubations), before being underlain with methyl salicylate. The brain was allowed to sink, before the methyl salicylate was refreshed (2 × 30 minute incubations).

### Confocal imaging

All imaging was performed on a confocal laser-scanning microscope (Leica TCS SP8, Leica Microsystem, Mannheim, Germany) using a 10× dry objective with a numerical aperture of 0.4 (Leica Material No. 11506511), a mechanical *z*-step of 2 μm and an *x-y* resolution of 512 × 512 pixels. Imaging the whole brain required capturing 3×2 tiled stacks in the *x-y* dimensions (20% overlap) that were automatically merged in Leica Applications Suite Advanced Fluorescence software. Each brain was scanned from the posterior and anterior side to span the full *z*-dimension of the brain. These image stacks were then merged in Amira 3D analysis software 5.5 (FEI Visualization Sciences Group; custom module ‘Advanced Merge’). The *z*-dimension was scaled 1.52× to correct the artifactual shortening associated with the 10× air objective (Heinze and Reppert, 2012; Montgomery and Ott, 2015). Images that illustrate key morphological details were captured separately as single confocal sections with an *x-y* resolution of 1024 × 1024 pixels.

### Neuropil segmentations and volumetric reconstructions

We assigned image regions to anatomical structures in the Amira 5.5 *labelfield* module by defining outlines based on the brightness of the synapsin immunofluorescence. Within each stack, every forth or fifth image was manually segmented and interpolated in the *z*-dimension across all images that contain the neuropil of interest. The *measure statistics* module was used to determine volumes (in μm^3^) for each neuropil. 3D polygonal surface models of the neuropils were constructed from the smoothed labelfield outlines (*SurfaceGen* module). The color code used for the neuropils in the 3D models is consistent with previous neuroanatomical studies of insect brains (Brandt et al., 2005; Kurylas et al., 2008; El Jundi et al., 2009a, b; Dreyer et al., 2010; Heinze and Reppert, 2012; Montgomery and Ott, 2015).

The whole-brain composite stacks were used to reconstruct and measure six paired neuropils in the optic lobes, and seven paired and two unpaired neuropils in the midbrain where distinct margins in staining intensity delineate their margins. All paired neuropils were measured on both sides of the brain in wild-caught individuals to permit tests of asymmetry, yielding two paired measurements per brain (*i.e. N* = 10 × 2) for each structure. We found no evidence of volumetric asymmetry for either species (p > 0.05 for each neuropil in a paired t-tests) and therefore summed the volumes of paired neuropil to calculate the total volume of that structure. In insectary-reared individuals we subsequently measured the volume of paired neuropil from one hemisphere, chosen at random, and multiplied the measured volume by two. We measured the total neuropil volume of the midbrain to permit statistical analyses that control for allometric scaling. In keeping with the earlier lepidopteran literature, we use the term ‘midbrain’ for the fused central mass that comprises the protocerebral neuromere excluding the optic lobes, the deuto- and tritocerebral neuromeres, and the sub-esophageal neuromeres. For the following statistical analyses we analyzed the central body as a single structure and, unless otherwise stated, summed the volumes of the mushroom body lobes and peduncles.

### Intraspecific statistical analyses

In all statistical analyses continuous variables were log_10_-transformed. Unpaired two-tailed two-sample *t-*tests were used to test for volumetric differences between sexes or groups. We found no evidence of sexual dimorphism in neuropil volume of wild caught individuals that could not be explained by allometric scaling and therefore combined male and female data.

All statistical analyses were performed in R v.3.1 (R Development Core Team, 2008). Our analyses focused on two intra-specific comparisons: i) we compared ‘young’ and ‘old’ insectary-reared individuals and interpret significant differences as evidence for post-eclosion growth; and ii) we compared wild-caught individuals with ‘old’ insectary-reared individuals and interpret significant differences as evidence for environmentally induced, or experience dependent plasticity. These comparisons were made by estimating the allometric relationship between each neuropil and a measure of overall brain size (total volume of the midbrain minus the combined volume of all segmented neuropil in the midbrain: ‘rest of midbrain’, rMid) using the standard allometric scaling relationship: log *y* = β log *x* + α. We used standard major axis regressions in the SMATR v.3.4-3 (Warton et al., 2012) to test for significant shifts in the allometric slope (β). Where we identified no heterogeneity in β we performed two further tests: 1) for differences in α that suggest discrete ‘grade-shifts’ in the relationship between two variables, 2) for major axis-shifts along a common slope. Patterns of brain:body allometry were explored in a similar manner, using total neuropil volume as the dependent variable (summed volumes of all optic lobes neuropil plus the total midbrain volume), and comparing the results obtained using alternative body size measurements as the independent variable. We also present the effect size, measured by the correlation coefficient (*r*). Effect sizes of 0.1<*r*<0.3 are interpreted as ‘small’ effects, 0.3<*r*<0.5 ‘medium’ effects, and *r*<0.5 ‘large’ effects (Cohen, 1988).

### Interspecific statistical analyses

To analyze interspecific patterns of divergence in brain composition we collected published data for neuropil volumes of four other Lepidoptera; *D. plexippus*; (Heinze and Reppert, 2012), *G. zavaleta* (Montgomery and Ott, 2015), *M. sexta*; (El Jundi et al., 2009a) and *H. virescens* (Kvello et al., 2009). Data were available for eight neuropils across all four species. Relative size was measured by calculating the residuals from a phylogenetically-corrected least squares (PGLS) linear regression between each structure and the rest of the brain (whole brain or midbrain as indicated) performed in BayesTraits (freely available from www.evolution.rdg.ac.uk; Pagel, 1999). For this analysis, a phylogeny of the six species was created using data on two loci, *COI* and *EF1a* (GenBank Accession IDs, *COI*: EU069042.1, GU365908.1, JQ569251.1, JN798958.1, JQ539220.1, HM416492.1; *EF1a*: EU069147.1, DQ157894.1, U20135.1, KC893204.1, AY748017.1, AY748000.1). The data were aligned and concatenated using MUSCLE (Edgar, 2004), before constructing a maximum likelihood tree in MEGA v.5 (Tamura et al., 2011). Differences in brain composition across species were analyzed by Principal Component analysis of these data, and visualized as biplots (Greenacre, 2010) in R package *ggbiplot* (V.Q. Vu, https://github.com/vqv/ggbiplot). Finally, we extended our phylogenetic analysis across insects using a similar approach. We restricted this analysis to volumetric data collected with similar methodology (Rein et al., 2002; Brandt et al., 2005; Kurylas et al., 2008; Dreyer et al., 2010; Ott and Rogers, 2010; Wei et al., 2010). The phylogenetic relationship of these insects was taken from Trautwein et al. (2012).

## RESULTS

### General layout of the *Heliconius* brain

The overall layout and morphology of the *Heliconius* brain (Fig. 1) is similar to that of other Lepidoptera (El Jundi et al., 2009; Kvello et al., 2009; Heinze and Reppert, 2012a; Montgomery and Ott, 2015). The midbrain forms a single medial mass, containing the supra-esophageal ganglion to which the sub-esophageal ganglion is fused. Together with the rest of the midbrain (rMid), which lacks distinct internal boundaries and was therefore unsuitable for further segmentation in the current analysis, we measured the volumes of six paired neuropils in the optic lobes, and eight paired and two unpaired neuropils in the midbrain in 59 individuals across both species (Table 1).

**Figure 1:**
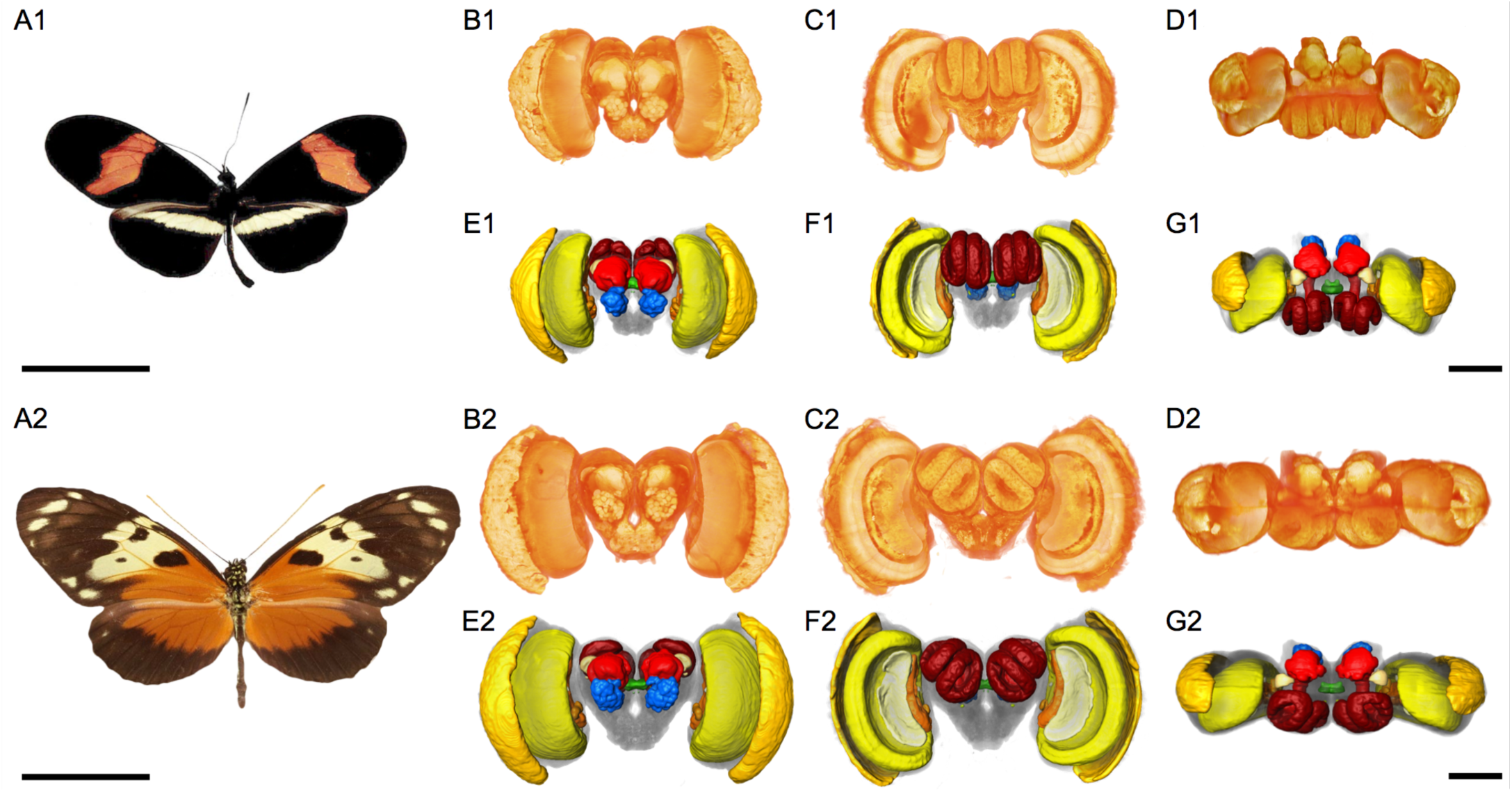
Overview of the anatomy of the Heliconius brain. 3D models of *H. erato* (A1–G1) and *H. hecale* (A2–G2). B1–D1 and B2–D2: Volume rendering of sin immunofluorescence showing the surface morphology of the brain neuropil from the ior (A1/A2), posterior (B1/B2), and dorsal (C1/C2) view. E1–G1 and E2–G2: Surface structions of the major neuropil compartments from the anterior (D1/D2), posterior (E1/E2), orsal (F1/F2) view. Neuropil in yellow-orange: visual neuropil, green: central complex, blue: nal lobes, red: mushroom bodies. See Figures 2–4 for further anatomical detail. The iduals displayed are male. Images in A1/A2 are from Warren et al. (2013). Scale bars = 25 mm in A1/A2; 500 μm in B1–D1/B2–D2.

**Table 1:**
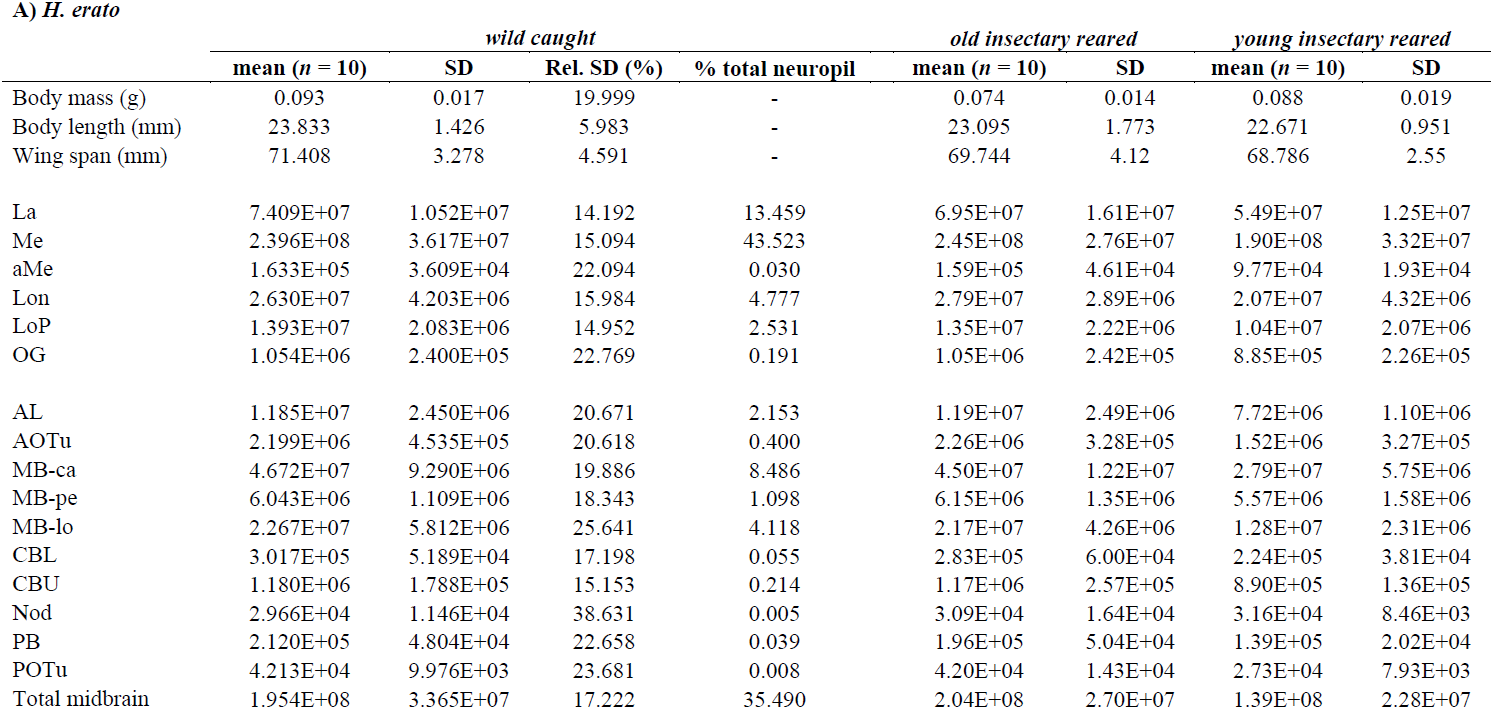
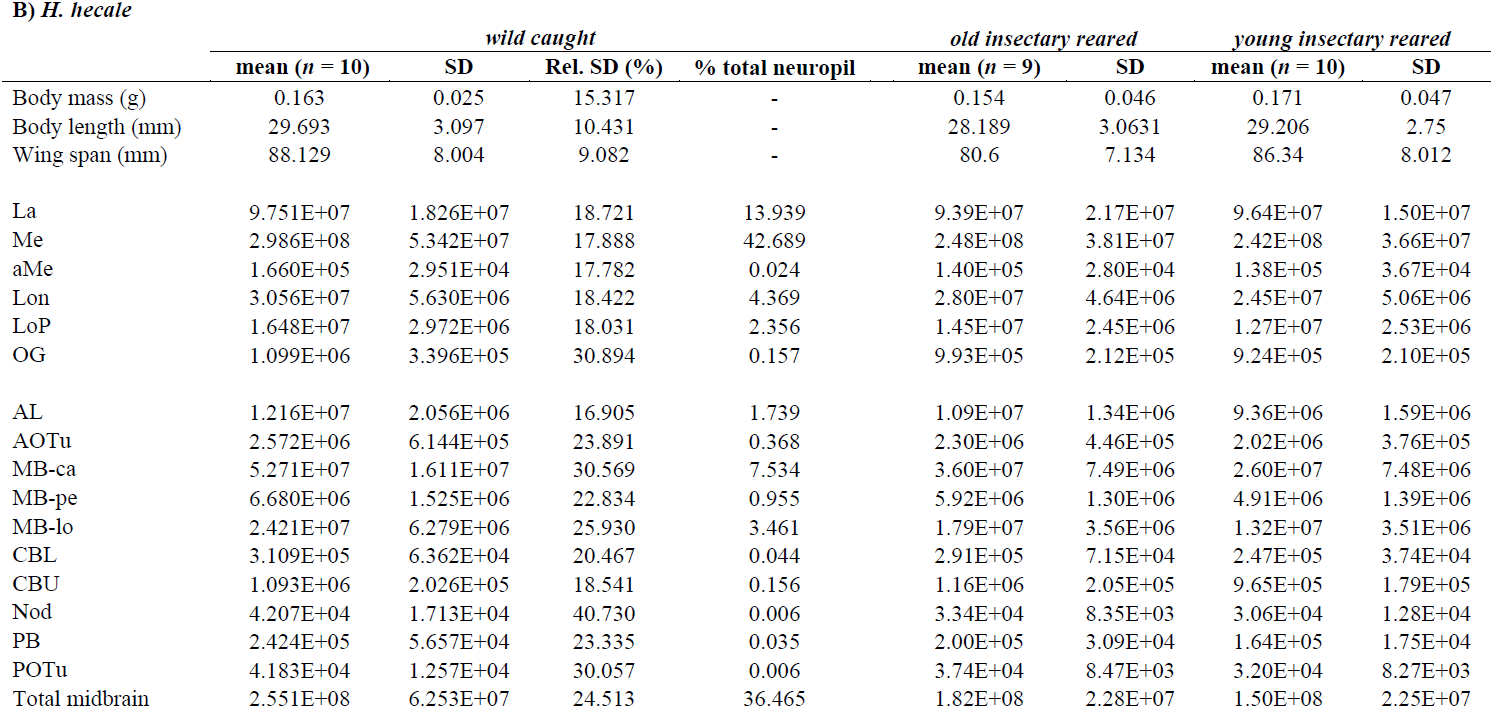
Neuropil volumes and body size of A) *H. erato* and B) *H. hecale*

|  | *wild caught* |  |  |  | *old insectary reared* |  | *young insectary reared* |  |
| --- | --- | --- | --- | --- | --- | --- | --- | --- |
|  | mean (*n* = 10) | SD | Rel. SD (%) | % total neuropil | mean (*n* = 10) | SD | mean (*n* = 10) | SD |
| Body mass (g) | 0.093 | 0.017 | 19.999 | - | 0.074 | 0.014 | 0.088 | 0.019 |
| Body length (mm) | 23.833 | 1.426 | 5.983 | - | 23.095 | 1.773 | 22.671 | 0.951 |
| Wing span (mm) | 71.408 | 3.278 | 4.591 | - | 69.744 | 4.12 | 68.786 | 2.55 |
| La | 7.409E+07 | 1.052E+07 | 14.192 | 13.459 | 6.95E+07 | 1.61E+07 | 5.49E+07 | 1.25E+07 |
| Me | 2.396E+08 | 3.617E+07 | 15.094 | 43.523 | 2.45E+08 | 2.76E+07 | 1.90E+08 | 3.32E+07 |
| aMe | 1.633E+05 | 3.609E+04 | 22.094 | 0.030 | 1.59E+05 | 4.61E+04 | 9.77E+04 | 1.93E+04 |
| Lon | 2.630E+07 | 4.203E+06 | 15.984 | 4.777 | 2.79E+07 | 2.89E+06 | 2.07E+07 | 4.32E+06 |
| LoP | 1.393E+07 | 2.083E+06 | 14.952 | 2.531 | 1.35E+07 | 2.22E+06 | 1.04E+07 | 2.07E+06 |
| OG | 1.054E+06 | 2.400E+05 | 22.769 | 0.191 | 1.05E+06 | 2.42E+05 | 8.85E+05 | 2.26E+05 |
| AL | 1.185E+07 | 2.450E+06 | 20.671 | 2.153 | 1.19E+07 | 2.49E+06 | 7.72E+06 | 1.10E+06 |
| AOTu | 2.199E+06 | 4.535E+05 | 20.618 | 0.400 | 2.26E+06 | 3.28E+05 | 1.52E+06 | 3.27E+05 |
| MB-ca | 4.672E+07 | 9.290E+06 | 19.886 | 8.486 | 4.50E+07 | 1.22E+07 | 2.79E+07 | 5.75E+06 |
| MB-pe | 6.043E+06 | 1.109E+06 | 18.343 | 1.098 | 6.15E+06 | 1.35E+06 | 5.57E+06 | 1.58E+06 |
| MB-lo | 2.267E+07 | 5.812E+06 | 25.641 | 4.118 | 2.17E+07 | 4.26E+06 | 1.28E+07 | 2.31E+06 |
| CBL | 3.017E+05 | 5.189E+04 | 17.198 | 0.055 | 2.83E+05 | 6.00E+04 | 2.24E+05 | 3.81E+04 |
| CBU | 1.180E+06 | 1.788E+05 | 15.153 | 0.214 | 1.17E+06 | 2.57E+05 | 8.90E+05 | 1.36E+05 |
| Nod | 2.966E+04 | 1.146E+04 | 38.631 | 0.005 | 3.09E+04 | 1.64E+04 | 3.16E+04 | 8.46E+03 |
| PB | 2.120E+05 | 4.804E+04 | 22.658 | 0.039 | 1.96E+05 | 5.04E+04 | 1.39E+05 | 2.02E+04 |
| POTu | 4.213E+04 | 9.976E+03 | 23.681 | 0.008 | 4.20E+04 | 1.43E+04 | 2.73E+04 | 7.93E+03 |
| Total midbrain | 1.954E+08 | 3.365E+07 | 17.222 | 35.490 | 2.04E+08 | 2.70E+07 | 1.39E+08 | 2.28E+07 |

**B) *H. hecale***
|  | *wild caught* |  |  |  | *old insectary reared* |  | *young insectary reared* |  |
| --- | --- | --- | --- | --- | --- | --- | --- | --- |
|  | mean (*n* = 10) | SD | Rel. SD (%) | % total neuropil | mean (*n* = 9) | SD | mean (*n* = 10) | SD |
| Body mass (g) | 0.163 | 0.025 | 15.317 | - | 0.154 | 0.046 | 0.171 | 0.047 |
| Body length (mm) | 29.693 | 3.097 | 10.431 | - | 28.189 | 3.0631 | 29.206 | 2.75 |
| Wing span (mm) | 88.129 | 8.004 | 9.082 | - | 80.6 | 7.134 | 86.34 | 8.012 |
| La | 9.751E+07 | 1.826E+07 | 18.721 | 13.939 | 9.39E+07 | 2.17E+07 | 9.64E+07 | 1.50E+07 |
| Me | 2.986E+08 | 5.342E+07 | 17.888 | 42.689 | 2.48E+08 | 3.81E+07 | 2.42E+08 | 3.66E+07 |
| aMe | 1.660E+05 | 2.951E+04 | 17.782 | 0.024 | 1.40E+05 | 2.80E+04 | 1.38E+05 | 3.67E+04 |
| Lon | 3.056E+07 | 5.630E+06 | 18.422 | 4.369 | 2.80E+07 | 4.64E+06 | 2.45E+07 | 5.06E+06 |
| LoP | 1.648E+07 | 2.972E+06 | 18.031 | 2.356 | 1.45E+07 | 2.45E+06 | 1.27E+07 | 2.53E+06 |
| OG | 1.099E+06 | 3.396E+05 | 30.894 | 0.157 | 9.93E+05 | 2.12E+05 | 9.24E+05 | 2.10E+05 |
| AL | 1.216E+07 | 2.056E+06 | 16.905 | 1.739 | 1.09E+07 | 1.34E+06 | 9.36E+06 | 1.59E+06 |
| AOTu | 2.572E+06 | 6.144E+05 | 23.891 | 0.368 | 2.30E+06 | 4.46E+05 | 2.02E+06 | 3.76E+05 |
| MB-ca | 5.271E+07 | 1.611E+07 | 30.569 | 7.534 | 3.60E+07 | 7.49E+06 | 2.60E+07 | 7.48E+06 |
| MB-pe | 6.680E+06 | 1.525E+06 | 22.834 | 0.955 | 5.92E+06 | 1.30E+06 | 4.91E+06 | 1.39E+06 |
| MB-lo | 2.421E+07 | 6.279E+06 | 25.930 | 3.461 | 1.79E+07 | 3.56E+06 | 1.32E+07 | 3.51E+06 |
| CBL | 3.109E+05 | 6.362E+04 | 20.467 | 0.044 | 2.91E+05 | 7.15E+04 | 2.47E+05 | 3.74E+04 |
| CBU | 1.093E+06 | 2.026E+05 | 18.541 | 0.156 | 1.16E+06 | 2.05E+05 | 9.65E+05 | 1.79E+05 |
| Nod | 4.207E+04 | 1.713E+04 | 40.730 | 0.006 | 3.34E+04 | 8.35E+03 | 3.06E+04 | 1.28E+04 |
| PB | 2.424E+05 | 5.657E+04 | 23.335 | 0.035 | 2.00E+05 | 3.09E+04 | 1.64E+05 | 1.75E+04 |
| POTu | 4.183E+04 | 1.257E+04 | 30.057 | 0.006 | 3.74E+04 | 8.47E+03 | 3.20E+04 | 8.27E+03 |
| Total midbrain | 2.551E+08 | 6.253E+07 | 24.513 | 36.465 | 1.82E+08 | 2.28E+07 | 1.50E+08 | 2.25E+07 |

### Sensory neuropil

The large optic lobes (OL; Fig. 2) account for approximately 64% of the total brain volume. As is the case in both *D. plexippus* and *G. zavaleta* the lamina (La), two-layered medulla (Me) (Fig. 2E), accessory medulla (aMe), lobula (Lob) and lobula plate (Lop) are well defined and positioned in the OL as nested structures from lateral to medial (Fig. 2A). The La has a distinct, brightly stained inner rim (iRim; Fig. 2E), a feature common to all diurnal butterflies analyzed thus far (Heinze and Reppert, 2012; Montgomery and Ott, 2015). In common with *D. plexippus* we identify a thin strip of irregularly shaped neuropil running ventrally from the aME to the Me (Fig. 2G–H).

**Figure 2:**
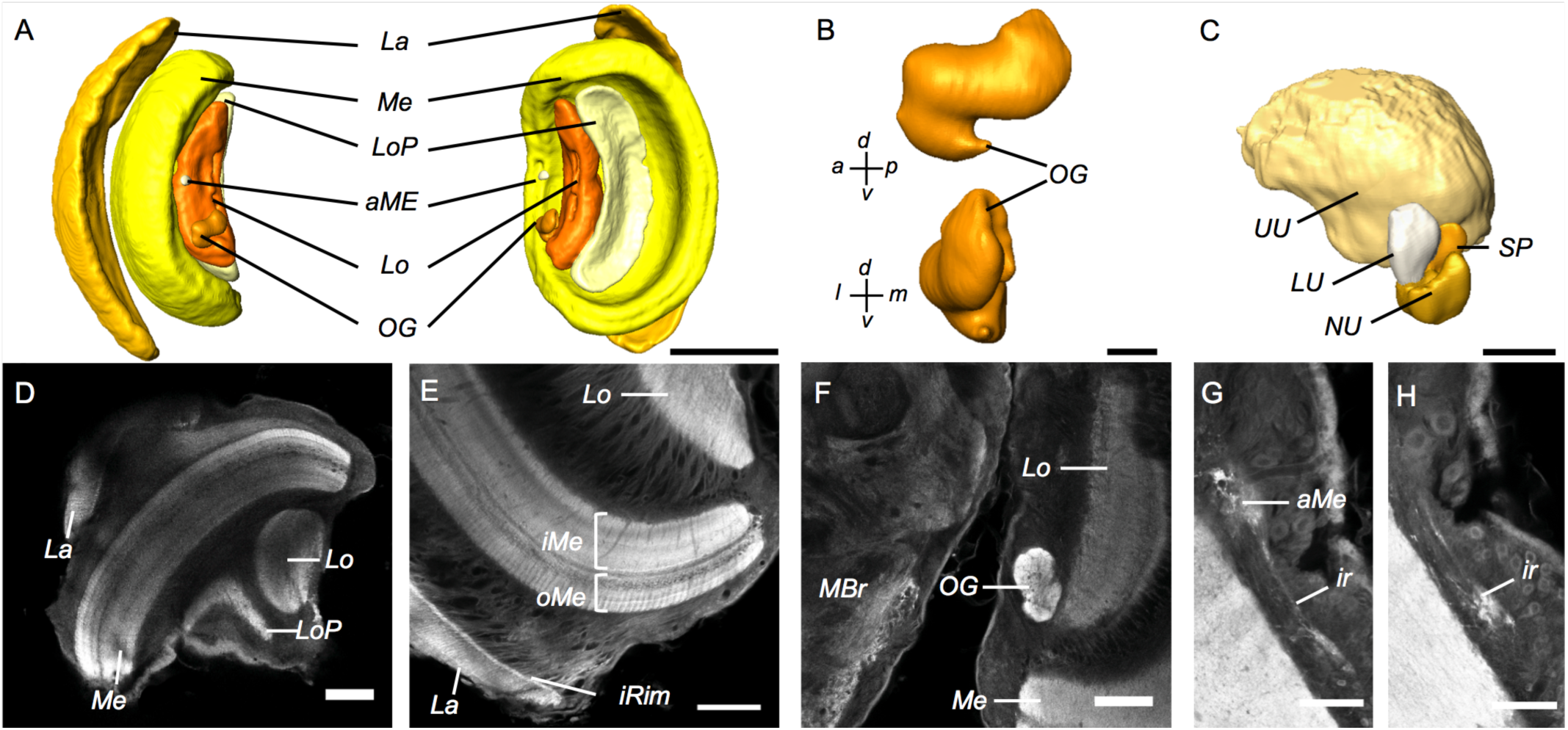
Anatomy of the major visual neuropils. A: Surface reconstructions of the optic lobe neuropils viewed from anterior (left image) and posterior image). They comprise the lamina (La), the medulla (Me), and accessory medulla (aMe), the (Lo), the lobula plate (LoP) and the optic glomerulus (OG). B: Surface reconstruction of the glomerulus (OG) viewed along the anterior-posterior axis (top) and an anterior view (bottom). Surface reconstruction of the anteriot optic tubercle (AOTu). D–J: Synapsin immunofluorescence gle confocal sections of the optic lobe of H. hecale. D: Horizontal section showing all four major optic lobe neuropils (La, Me, Lo, LoP). E: Frontal section showing the inner rim (iRim) of the lamina, a thin layer on its inner surface that is defined by intense synapsin immunofluorescence. Synapsin immunostaining also reveals the laminated structure of the medulla with two main subdivisions, the outer and inner medulla (oMe, iMe). F: The OG is located medially to the Lo; frontal section, the midbrain (MBr) occupies the left half of the frame. G,H: Frontal sections showing a small, irregular neuropil (ir) observed running from the anterior-ventral boundary of the aME as in *xippus* (Heinze and Reppert, 2012). All images are from male *H. hecale*.

We also identify a sixth neuropil in the OL that we believe to be homologous to the optic glomerulus (OG; Fig. 2B,F) identified in *D. plexippus* (Heinze and Reppert, 2012), which is absent in other lepidopteran brains described to date and was postulated to be Monarch-specific. As in *D. plexippus* this neuropil is a multi-lobed, irregularly shaped structure positioned to the medial margin of the Lob with which it appears to be connected. In *Heliconius* the OG is not as extended in the anterior margin as in *D. plexippus* and is subsequently confined to the OL, without protrusion into the optic stalk or midbrain (Fig. 2A,B,F). The position of the OG in *Heliconius* is also similar to that of a much smaller neuropil observed in *G. zavaleta* (Montgomery and Ott, 2015) that may be homologous.

The midbrain contains further neuropils with primary functions in processing visual information that include the anterior optic tubercule (AOTu). We identify the same four components of the AOTu previously described in *D. plexippus* and *G. zavaleta* butterflies (Heinze and Reppert, 2012; Montgomery and Ott, 2015): the small, closely clustered nodular unit (NU), strap (SP) and lower unit (LU), and the much larger upper unit (UU) (Fig. 2C). As in other butterflies, the UU is expanded compared with nocturnal moths (El Jundi et al., 2009; Kvello et al., 2009). The proportion of total neuropil comprised of the AOTu is, however, larger in *D. plexippus* (0.74%) than *Heliconius* (0.40% in *H. hecale* and 0.37% in *H. erato*).

The antennal lobes (AL), the primary olfactory neuropil, are comprised of small, round glomeruli that are innervated by axons from olfactory receptor neurons in the antennae. These glomeruli are arranged around a Central Fibrous Neuropil (CFN) (Figure 3A,B). In *Heliconius* the AL comprises 2% of the total brain neuropil volume, and contains approximately 68 glomeruli (estimated in one individual of each sex: *H. erato* ♂ = 69, ♀ = 68; *H. hecale* ♂ = 68, ♀ = 67) which matches closely the number of olfactory receptor genes (70) identified in the *H. melpomene* genome (Dasmahapatra et al., 2012). We found no expanded macro-glomeruli complex (MGC) or obvious candidates for sexually dimorphic glomeruli. This is in keeping with all diurnal butterflies described to date (Rospars, 1983; Heinze and Reppert, 2012; Carlsson et al., 2013), with the exception of the more olfactorily orientated *G. zavaleta* (Montgomery and Ott, 2015).

**Figure 3:**
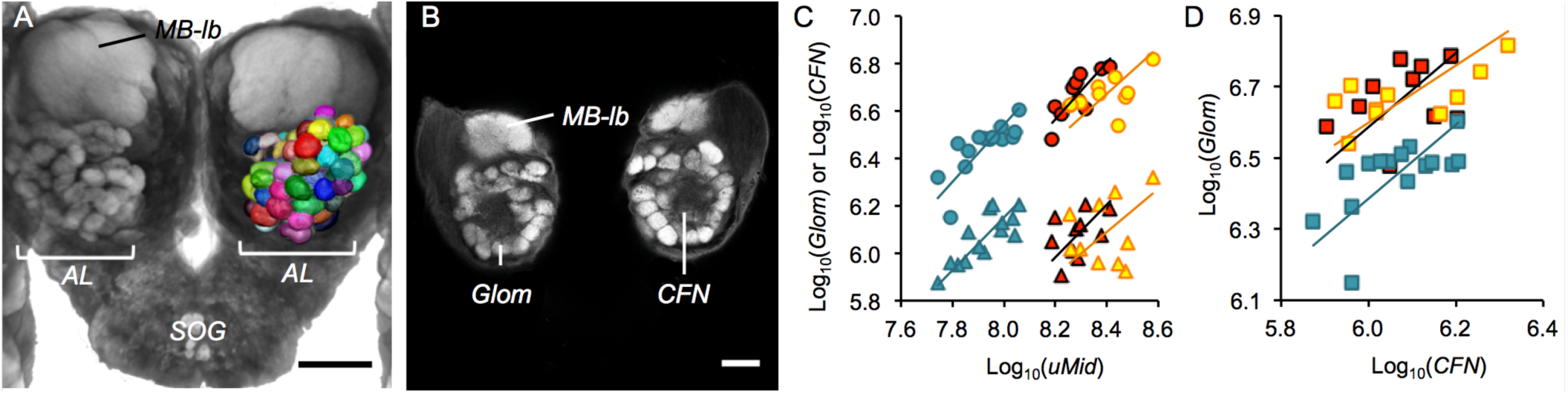
Anatomy of the antennal lobe. A: 3D reconstruction of individual antennal lobe (AL) glomeruli superimposed on a volume rendering of the anterior surface of the midbrain. B: Synapsin immunofluorescence in a single frontal confocal section showing the glomeruli (Glom) surrounding the central fibrous neuropil (CFN). A–B are from male *H. hecale*. C,D: Allometric grade-shifts between Glomerular (circles) or triangles) volume and unsegmented midbrain volume (C), and between Glomerular and CFN (D) in *G. zavaleta* (solid blue), *H. erato* (black filled with red) and *H. hecale* (orange filled with yellow). Scale bars = 500 μm in A; 50 μm in B,C,G,H; 100 μm in B–F, J; 200 μm in I.

We took advantage of comparable datasets for *H. erato, H. hecale* and *G. zavaleta* to investigate whether changes in relative AL volume are due to an increased volume of glomeruli or CFN. Both glomerular and CFN volume are larger in *G. zavaleta* relative to midbrain volume, as indicated by significant grade-shifts in allometric scaling in *G. zavaleta* and *Heliconius* (glomerular, *H. erato*: Wald χ2 = 10.709, p = 0.001; *H. hecale*: Wald χ2 = 9.139, p = 0.003; CFN, *H. erato*: Wald χ2 = 30.282, p < 0.001; *H. hecale*: Wald χ2 = 26.638, p < 0.001). However, CFN expansion in *G. zavaleta* is disproportionately large, driving a grade-shift in the scaling relationship between glomerular and CFN volume in *G. zavaleta* when compared with either *Heliconius* (*H. erato*: Wald χ2 = 19.680, p < 0.001; *H. hecale*: Wald χ2 = 31.663, p < 0.001; Fig. 3D).

### Central complex

The central complex is a multimodal integration center linked to a range of functions from locomotor control to memory (Pfeiffer and Homberg, 2014). Within the limitations of the current analysis, the anatomy of the *Heliconius* central complex shows strong conservation with *D. plexippus* and *G. zavaleta* (Heinze and Reppert, 2012; Montgomery and Ott, 2015). The central body (CB) is positioned along the midline of the central brain and is formed of two neuropils, the upper (CBU) and lower (CBL) divisions, which are associated with small paired neuropils, the noduli (No), located ventrally to the CB (Fig. 4A–D,G). Two further paired neuropils, the protocerebral bridge (PB; Fig. 4A,E) and posterior optic tubercles (POTu; Fig. 4A,F), are positioned towards the posterior margin of the brain.

**Figure 4:**
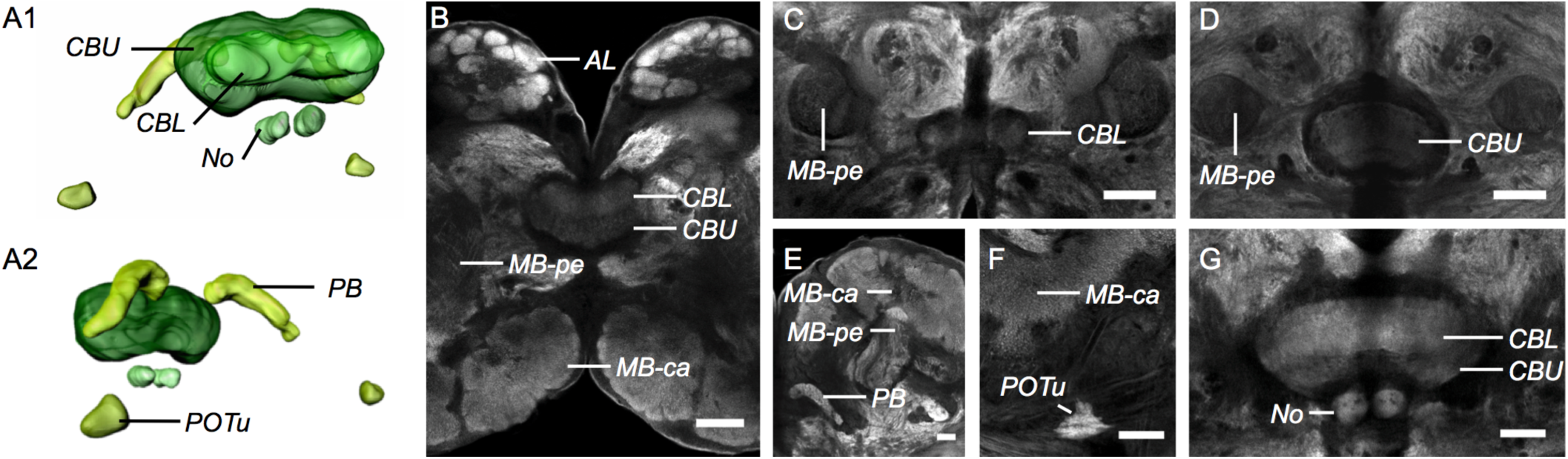
Anatomy of the central complex. A1/A2: Surface reconstruction of the central complex from an anterolateral (A1) and oblique roventral (A2) view, showing the upper and lower subunit of the central body (CBU, CBL), the (No), the protocerebral bridge (PB) and posterior optic tubercles (POTu). B–G: Synapsin immunofluorescence in single confocal sections. B: Horizontal section showing the upper and lower it of the CB in relation to the antennal lobes (AL) and the calyx (MB-ca) and pedunculus (MB-pe) of the mushroom body. C,D: Frontal confocal sections at the level of the CBL (C) and CBU (D); CB subunits are flanked by the profiles of the vertically running MB-pe on either side. E: Frontal section showing the location of the PB ventrally to the MB-ca. F: POTu positioned ventrally to the MB-ca in a frontal section. G: Frontal section showing position of the paired No ventrally to CBL and CBU. All images are from a male H. hecale. Scale bars = 100 μm in B–D, G; 50 μm in E,F.

### Mushroom bodies

The most striking aspect of *Heliconius* brain morphology is the hugely expanded mushroom bodies which span the depth of the brain along the anterior-posterior axis (Fig. 5). On the anterior side, the mushroom body lobes (MB-lo) lie above the AL. As in *D. plexippus* (Heinze and Reppert, 2012) the distinct lobe structure observed in moths (El Jundi et al., 2009; Kvello et al., 2009) is lost, possibly due to extensive expansion. The only identifiable feature is a lobe curving round the medial margin, likely to be part of the vertical lobe (Fig. 5D,F). The MB-lo merges with the cylindrical pedunculus (MB-pe) that extends to the posterior midbrain. The boundary between the MB-lo and MB-pe is not distinct. The combined volume of the MBlo+pe accounts for 12.2% of total midbrain volume in *H. hecale* and 14.6% of total midbrain volume in *H. erato*, at least twice that reported for other Lepidoptera (Sjöholm et al., 2005; El Jundi et al., 2009; Kvello et al., 2009; Heinze and Reppert, 2012a; Montgomery and Ott, 2015). At the posterior end, the MB-pe splits into two roots that are encircled by the mushroom body calyx (MB-ca; Fig. 5A,H,K). A Y-tract runs parallel to the MB-pe from the posterior boundary of the MB-lo to the junction between the MB-pe and MB-ca. The Y-tract ventral loblets seen in other Lepidoptera (El Jundi et al., 2009; Kvello et al., 2009) are not distinct, having merged with the MB-lo (Fig. 5A,J,N).

**Figure 5:**
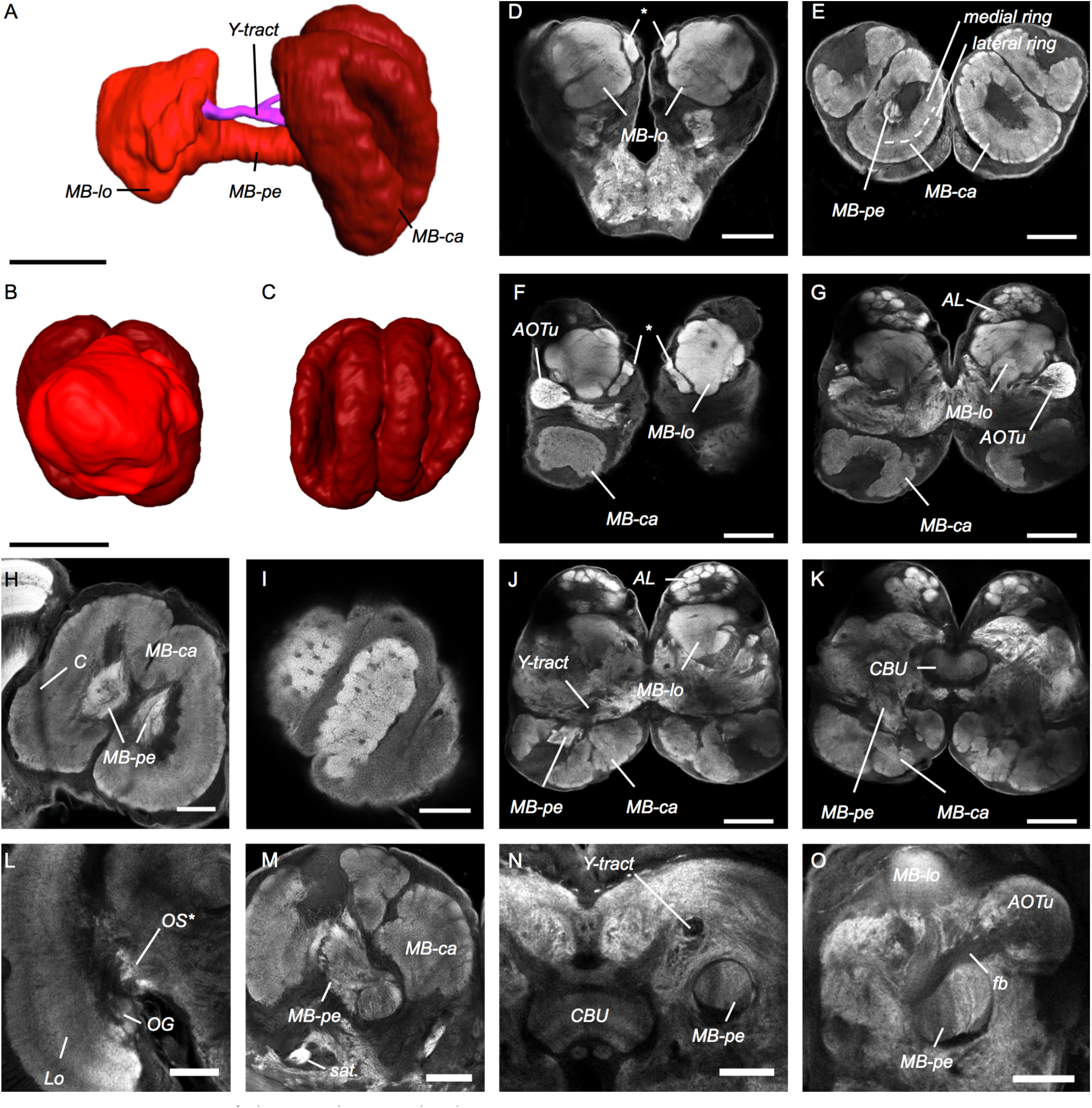
Anatomy of the mushroom body. A–C: Surface reconstruction of the mushroom body viewed orthogonal to the anterior-posterior axis medial vantage point level with the peduncle (A); from anterior (B); and from posterior (C). The main components are the calyx (MB-ca) shown in dark red, and the peduncule (MB-pe) and lobes (MB-lo) shown in bright red. A Y-tract, shown in magenta, runs parallel and slightly medial to the MB-pe. D–O: Synapsin immunofluorescence in individual confocal sections. D: anterior view of the midbrain showing the MB-lo, an asterik indicates what is most likely the ventral lobe, otherwise the individual lobes and loblets of the MB-lo are fused. E: Frontal section at a posterior level near the end of the MB-pe, showing the profiles of the MB-ca with its zonation into an outer and a medial ring. F,G and J,K: Horizontal confocal sections through the midbrain at increasing depths from dorsal towards ventral, showing MB structure in relation to neighboring neuropil: the anterior optic tubercle (AOTu in F,G); the antennal lobe (AL in G,J); and the central body upper division (CBU in K). H: An example of a female *H. erato* where the MB-ca is deformed due expansion into the optic lobe and constriction (labeled C) at the optic stalk by the neural sheath surrounding the brain. I: Pitted surface of the MB-ca in a very posterior tangential horizontal section. The pitting is related to what appear to be columnar domains within the calyx neuropil (cf. MB-ca in J,K,M). L: Areas of intense synapsin staining in the optic stalk (OS*); Lo, lobula; OG, optic glomerulus. M: Frontal section near the base of the calyx (MB-ca) showing a satellite neuropil (sat.) located near to the MB-pe. N: A Y-tract runs parallel with, and dorsally and slightly medially to the MB-pe; both are seen in profile in this frontal section. O: A fiber bundle (fb) connected to the AOTu running near the junction between the MB-pe and MB-lo. With the exception of I, all images are from a male *H. hecale*. Scale bars A-G, J-K = 200 μm, H-I, L-O = 100 μm.

*Heliconius* have an un-fused double MB-ca with a deeply cupped morphology (Fig. 5A,C). Two concentric zones can be identified (Fig. 5E), though the boundary is not distinct throughout the depth of the neuropil. The MB-ca comprises 20.7% and 23.9% of total midbrain volume in *H. hecale* and *H. erato* respectively, at least three times greater than reported in other Lepidoptera (Sjöholm et al., 2005; El Jundi et al., 2009; Kvello et al., 2009; Heinze and Reppert, 2012a; Montgomery and Ott, 2015). In some individuals the MB-ca is so large that it protrudes into the OL resulting in a distortion of shape caused by constriction around the optic stalk (Fig. 5H). We also observe some degree of pitting in the posterior surface of the MB-ca (Fig. 5I). This pitting is related to radially arranged columnar domains that are apparent within the calycal neuropil (Fig. 5J,K). We do not observe any structure clearly identifiable as an accessory calyx. We do see a brightly stained globular neuropil below the MB-ca/pe junction but it is quite some distance away from the junction and lacks the ‘spotty’ appearance of the accessory calyx in *D. plexippus* (Heinze and Reppert, 2012). It seems more likely that this structure is a ‘satellite’ neuropil that is not part of the MB (Farris, 2005). Its position corresponds roughly to the medial end of the expanded OG in *D. plexippus.* In some preparations one can follow a narrow faint fiber tract from here to an area of more intense staining in the optic stalk and on to the medial margin of the OG. If this is a functional connection, it is conceivable that the medial expansion of the OG in *D. plexippus* occurred along this pre-existing pathway.

### Interspecific divergence in brain composition and mushroom body expansion in *Heliconius*

After correcting for allometric scaling by phylogenetically-corrected regressions against total neuropil volume, the six lepidopteran species can be separated along the first two principal components that together explain 90.7% of variance. PC1 (65.9% of Var) is heavily loaded by sensory neuropil in one direction, and the MB-ca and MB-lo+ped in the other (Table 2). PC2 (24.8% of Var) is heavily loaded by the Me in one direction and the AL and CB in the other. This roughly separates the six species into three pairs, representing (i) *H. hecale* and *H. erato*; (ii) the other diurnal butterflies, *D. plexippus* and *G. zavaleta*; and (iii) the night-flying moths, *H. virescens* and *M. sexta* (Fig. 6B). When midbrain neuropil are analyzed separately, PC1 (68.7% of Var) marks an axis dominated by AL, CB and MB, and PC2 (23.3% of Var) is strongly loaded by the AOTu (Fig. 6C). This leads to two clusters grouping (i) *H. hecale* and *H. erato*, which invest heavily in the mushroom body neuropil, and (ii) the night-flying moths and *G. zavaleta*, which invest heavily in olfactory neuropil; leaving *D. plexippus* isolated by its large AOTu volume.

**Table 2:** Loadings on Principal Components Analysis of the relative size of brain components across six Lepidoptera.

| **Neuropil** | **Loadings** |  |
| --- | --- | --- |
|  | **Residuals** |  |
|  | **PC1** | **PC2** |
| AL | -0.981 | -0.045 |
| CB L+U | -0.798 | 0.406 |
| MB-ca | 0.962 | 0.110 |
| MB-lo+pe | 0.952 | 0.231 |
| AOTu | -0.047 | 0.966 |

**B) Whole neuropil**
| **Neuropil** | **Loadings** |  |
| --- | --- | --- |
|  | **Residuals** |  |
|  | **PC1** | **PC2** |
| AL | 0.761 | 0.619 |
| CB L+U | 0.671 | 0.670 |
| MB-ca | -0.961 | 0.212 |
| MB-lo+pe | -0.942 | 0.222 |
| AOTu | 0.811 | 0.024 |
| Me | 0.042 | -0.949 |
| Lob | 0.920 | -0.354 |
| LoP | 0.962 | -0.167 |

**Figure 6:**
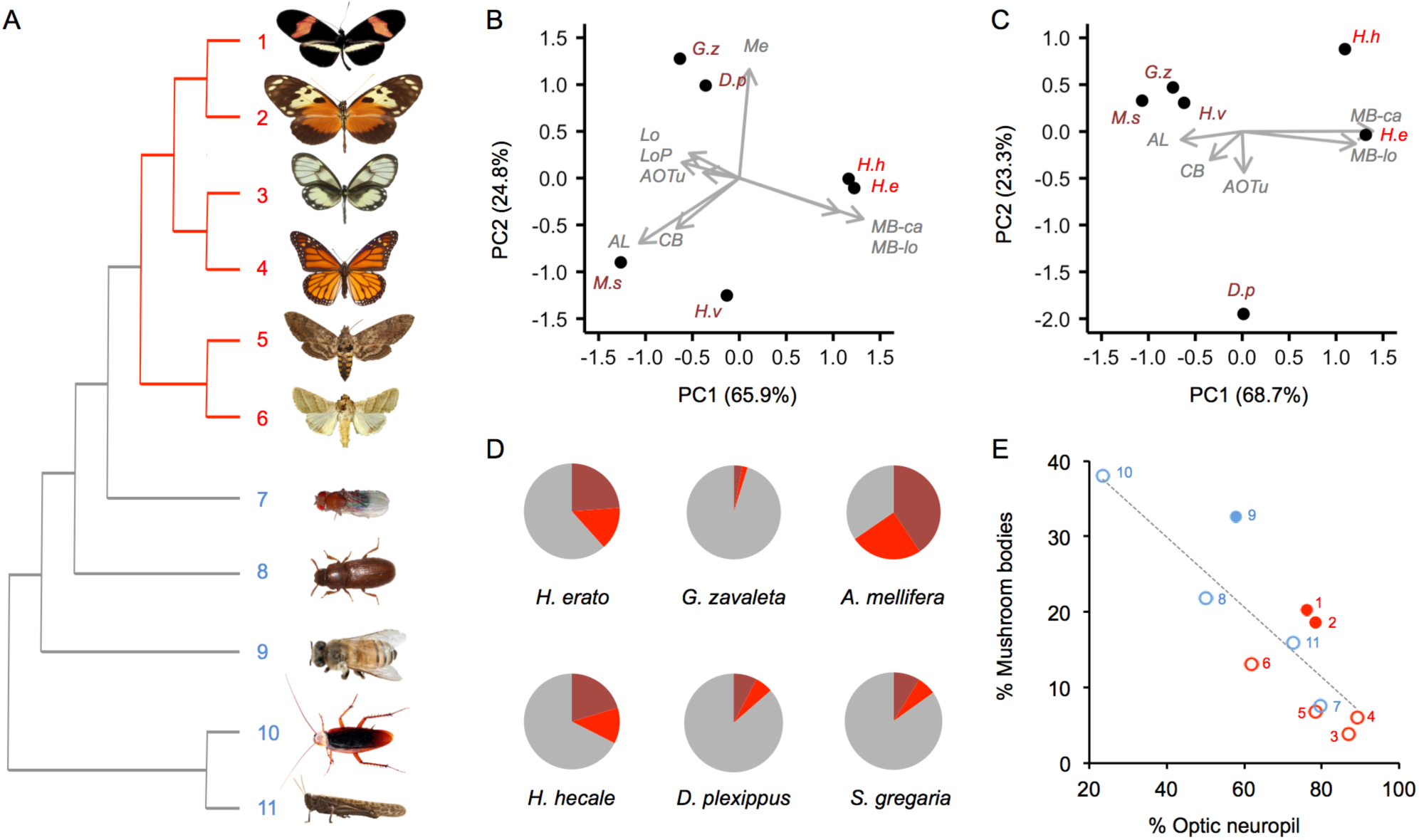
Divergence in brain structure across Lepidoptera, and in mushroom body size across insects. A: Phylogenetic relationships of Lepidoptera (red branches) and other insects (grey branches) for which directly comparable data are available. Branches are not drawn proportional to divergence dates, numbers refer to labels in panel E. B,C: Principal Component analysis of segmented neuropil volumes, corrected for allometric scaling with the unsegmented midbrain and phylogeny. B shows the results of an analysis using all neuropil. C shows the results of an analysis excluding the optic lobe neuropil. Species data points are indicated by the first letter of their genus and species name: D.p = *Danaus plexippus*; H.e = *Heliconius erato*; H.h = *H. hecale*; G.z = *Godyris zavaleta*; H.v = *Heliothis virescens*; M.s = *Manduca sexta*. D: The proportion of the midbrain occupied by MB-ca (dark red) and MB-lo+pe (light red) in four butterflies, and two other insects with fully comparable data. E: Across a wider sample of insects (shown in A), when expressed as a percentage of total volume of OL, AL, CB and MB, *Apis mellifera* (solid blue) and *Heliconius* (solid red) stand out as having expanded mushroom bodies, correcting for the size of the optic neuropil, compared to other Lepidoptera (unfilled red circles) and other insects (unfilled blue circles). The line was fitted by PGLS. All insect images in A are from Wikimedia commons and were released under the Creative Commons License, except *Heliconius* (see Fig. 1).

The combined volume of the calyx, pedunculus and lobes account for 13.7% of total brain neuropil volume in *H. erato*, and 11.9% in *H. hecale*. This is much larger than reported for any other Lepidoptera measured with similar methods (range 2.3–5.1%). Expressed as a percentage of the midbrain, to remove the effects of variation in the large OL, *H. erato* (38.5%) and *H. hecale* (32.9%) again exceed other Lepidoptera (4.8–13.5%) by 3–7 fold. These figures are also much larger than reported for *H. charithonia* (4.2% of total brain size) by Sivinski (1989), whose figures for other Lepidoptera are also much lower suggesting the difference is explained by variation in methodology.

Beyond Lepidoptera, the most comparable data available are from *Apis mellifera* (Brandt et al., 2005) and *Schistocerca gregaria* (Kurylas et al., 2008) for which mushroom body volume and midbrain volume are reported (Fig. 6D). In terms of raw volume (Table 1) *Heliconius* mushroom bodies are roughly equal in size to *A. mellifera*. However, in *A. mellifera* the mushroom bodies comprise 65.4% of the midbrain, (40.6% MB-ca, 24.8% MB-lo+ped) (Brandt et al., 2005), in gregarious-phase *S. gregaria* they comprise 15.1% (8.2% MB-ca including the accessory calyx, 6.3% MB-lo+ped) (Kurylas et al., 2008). Further comparisons can be made expressing mushroom body size as a percentage of segmented neuropil (Me+Lobula system, CB, MB and AL) that were labeled across a wider range species. As a ratio of percentage mushroom body volume to the percentage of the two other midbrain neuropil (AL and CB), *Heliconius* have ratios (*H. erato*: 6.4; *H. hecale*: 6.7) that exceed even *A. mellifera* (3.8). To account of the dominant effect of OL size on scaling with overall brain size we also analysed residual variance from a PGLS regression (Fig. 6E) between percentage OL and percentage MB volume. This shows *Heliconius* (*H. erato* +8.2; *H. hecale* +7.5) have the second largest residual MB size following *A. mellifera* (+11.9).

### Brain : body allometry

In both species, larger wild individuals have larger brains when using total neuropil volume and either body length or wingspan as measures of brain and body size (log-log SMA regression, *H. hecale,* body length *p* = 0.020; wingspan *p* = 0.019; *H. erato,* body length *p* = 0.011; wingspan *p* = 0.010). The brain size : body *mass* relationship is not significant in wild individuals (*H. hecale, p* = 0.055; *H. erato, p* = 0.863), most likely because body mass varies much with reproductive state and feeding condition. We therefore used body length as a proxy for body size to analyze the effect of age and experience on the relative size of the brain.

Both species showed a clear grade-shift with age towards increased relative brain size (*H. hecale:* Wald χ^2^ = 5.780, *p* = 0.016; *H. erato:* Wald χ^2^ = 10.124, *p* = 0.001). Body length was very similar in old and young individuals (*H. hecale t*_18_ = - 0.918, *p* = 0.371; *H. erato t*_17_ = 0.581, *p* = 0.568) suggesting the effect reflects an increase in absolute neuropil volume. Indeed, old individuals had significantly larger absolute midbrain volumes in both species (*H. erato: t*_17_ = 4.192, *p* = 0.001, *r* = 0.713; *H. hecale: t*_18_ = 3.054, *p* = 0.007, *r* = 0.595; Fig. 7A,D). An absolute increase in OL and total brain volume, however, was strongly supported only in *H. erato* (OL: *t*_17_ = 5.076, *p* < 0.001, *r* = 0.776; total, *t*_17_ = 5.153, *p* < 0.001, *r* = 0.708) and not evident in *H. hecale* (OL, *t*_18_ = 0.280, *p* = 0.783; total, *t*_18_ = 1.082, *p* = 0.293).

**Figure 7:**
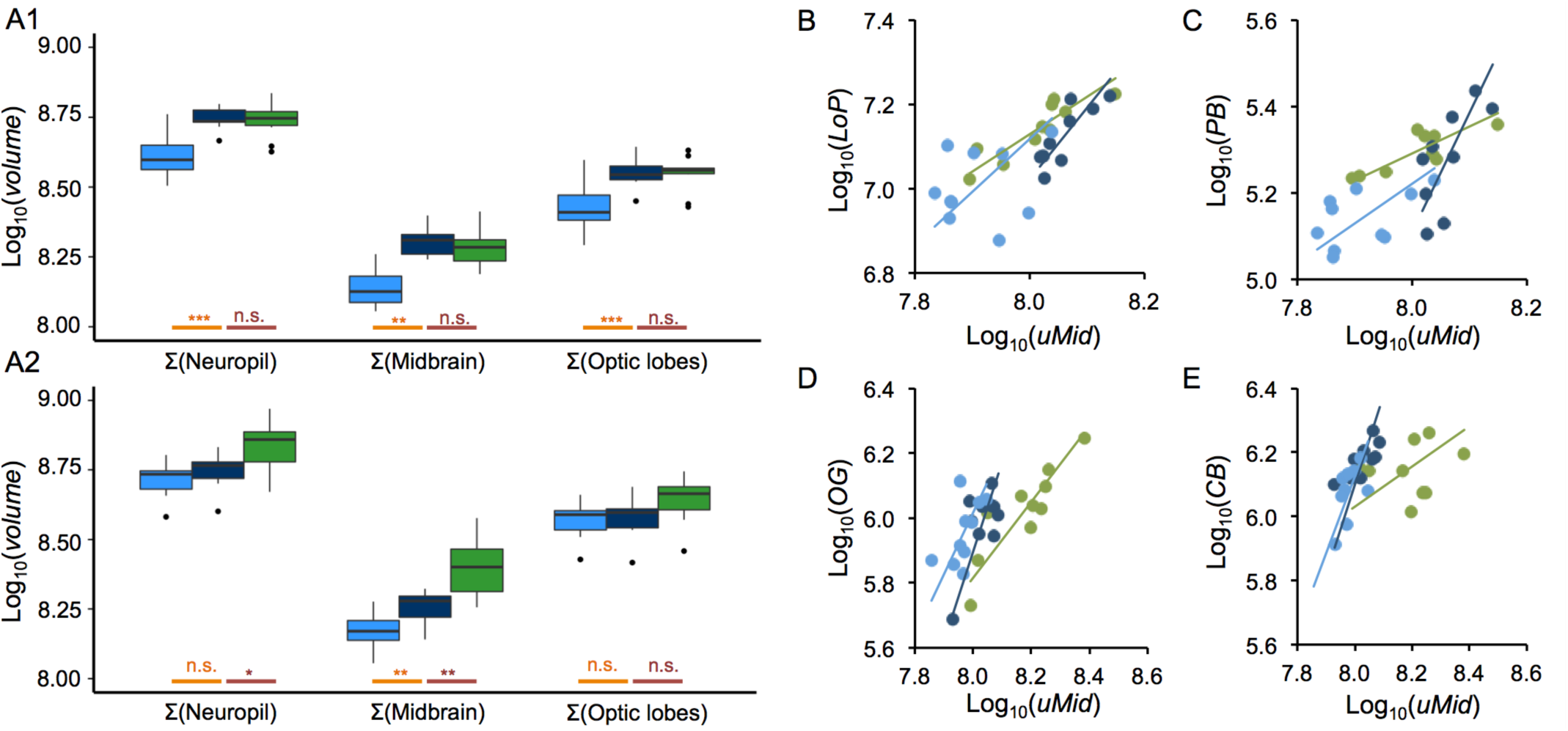
Age and environment dependent growth of brain components. A1,A2: Comparisons of raw volumes of total neuropil, total OL neuropil, total midbrain neuropil between wild-caught, old and young insectary-reared individuals of *H. erato* (A1) and *H. hecale* (A2). Significance of pair-wise comparisons are shown along the x-axis (young-old = orange; old-wild = dark red; n.s. = p>0.05, * = p<0.05, ** = p <0.01, *** = p < 0.001). B: Allometric scaling of LoP in *H. erato*. C: Allometric scaling of PB in *H. erato*. D: Allometric scaling of OG in *H. hecale*. E: Allometric scaling of CB in *H. hecale*. Note in E and F the shifts in allometry occur along the x-axis, this is explained by the large difference in unsegmented midbrain volume observed between wild-caught and old insectary-reared individuals in *H. hecale* as displayed in D.

Only *H. hecale* showed a clear response in overall brain size to experience. The total neuropil was 40% larger in wild-caught than in old insectary-reared individuals (*t*_17_ = 2.553, *p* = 0.020, *r* = 0.526) driven by a significant difference in midbrain volume (*t*_17_ = 3.658, *p* = 0.002, *r* = 0.664), but not OL volume (*t*_18_ = 1.728, *p* = 0.101; Fig. 7D). Although there was no matching difference in body length (*t*_18_ = 0.983, *p* = 0.436), a grade-shift towards larger relative brain size in wild *hecale* was not supported (Wald χ^2^ = 2.058, *p* = 0.151). However, we do observe a grade-shift when the midbrain is analyzed separately (Wald χ^2^ = 4.725, *p* = 0.030). No significant brain or body size differences were found between wild and old insectary-reared individuals in *H. erato* (total neuropil: *t*_17_ = -0.432, *p* = 0.671; midbrain: *t*_17_ = -0.732, *p* =0.474; OL: *t*_17_ = -0.123, *p* = 0.904; body length: *t*_17_ = 1.009, *p* = 0.327; Fig. 7A).

### Post-eclosion growth in the volume of individual neuropil regions

The age-related increase in overall absolute brain size in *H. erato* was reflected in volumetric increases in nearly all brain regions, with only the OG failing to show a significant expansion in old individuals (Table 3A). There was some evidence for age-related differences in the allometric scaling coefficients for aMe and PB, and for grade-shifts in OG and POTu, but these were weak relative to the strong major axis shifts observed for all neuropils investigated (Table 3A). The largest shifts were observed for the POTu (difference in fitted-axis mean, Δ_FA_ = 0.604), aME (Δ_FA_ = 0.536), MB-ca (Δ_FA_ = 0.496) and MB-lo+ped (Δ_FA_ = 0.393; Fig. 8A-C).

**Figure 8:**
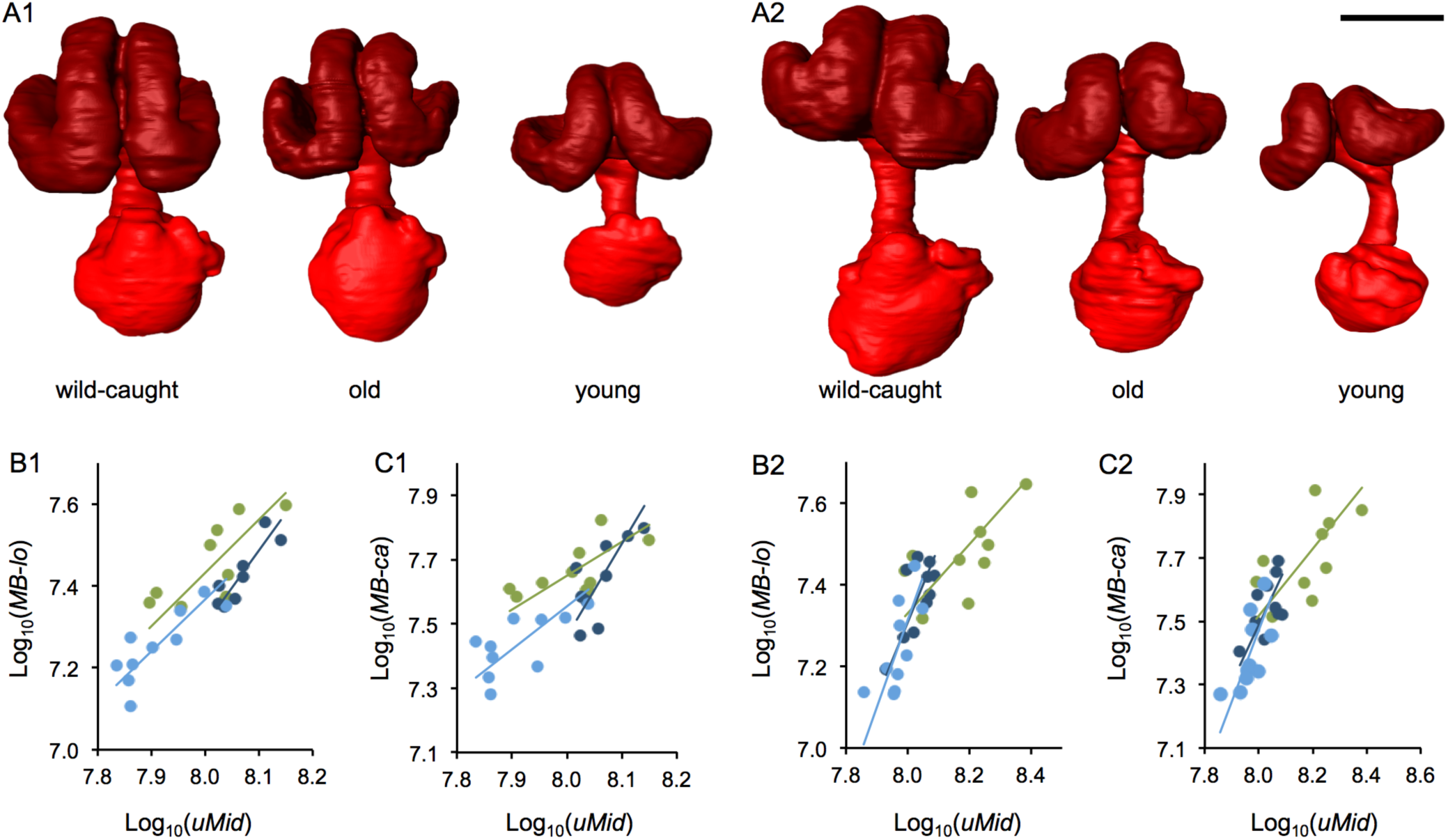
Age and environment dependent growth of the mushroom bodies. Surface reconstruction of the mushroom body viewed along the anterior-posterior axis for wild-caught, old and young insectary-reared individuals of H. erato (A1) and H. hecale (A2). Representative individuals were chosen as those closest to the group mean volume. Scale bar = 200 μm. B1-C1/B2-C2: allometric relationships between MB-lo+pe (B1/B2), or MB-ca (C1/C2),and the volume of the unsegmented midbrain (rMid) for *H. erato* (B1/C1) and *H. hecale* (B2/C2). Data for wild caught individuals are in green, data for old insectary-reared individuals in dark blue, and data for young insectary-reared individuals are in light blue. Allometric slopes for each group are shown, the slope, intercepts and major-axis means are compared in Table 3,4.

**Table 3:**
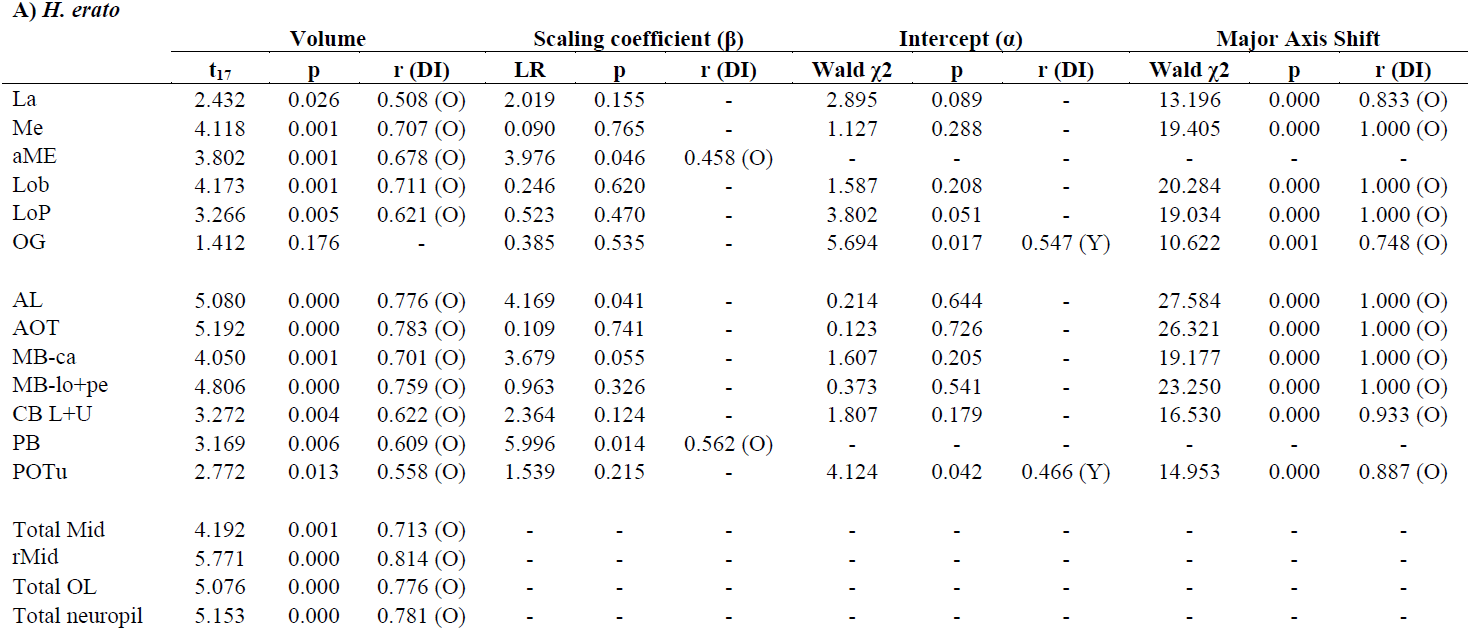
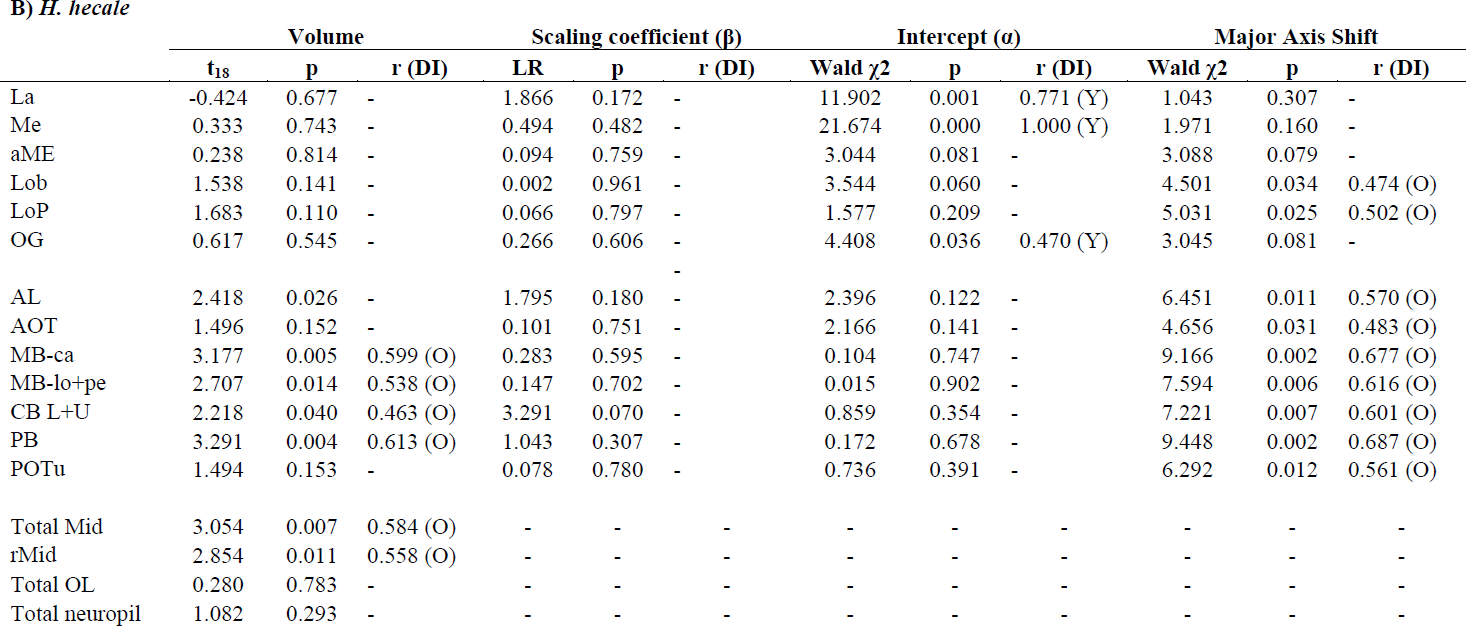
Comparisons between old (O) and young (Y) insectary-reared individuals for A) *H. erato* and B) *H. hecale. r* is the effect size. DI indicates the group with a higher value of α, β or fitted axis mean.

In contrast, in *H. hecale,* age-related size increases in volume were confined to the midbrain and not all segmented midbrain regions showed the same pattern of expansion; the rMid, components of the mushroom body complex, central complex and AL were all significantly larger in old individuals, but the AOTu, POTu and all optic lobe neuropil were not (Table 3B). Neuropil expansion appears to occur in a co-ordinated manner, such that the allometric relationship between each neuropil and rMid is maintained (Table 3B). The only exceptions were the La, Me and OG, which showed significant grade-shifts towards a reduced volume relative to rMid in old individuals. All other segmented neuropil showed major-axis shifts along a common slope towards higher values in old individuals (Table 3B). The largest shifts were observed in the mushroom body (MB-ca, Δ_FA_ = 0.279; MB-lo+ped, Δ_FA_ = 0.250; Fig. 8A1–C1).

### Experience-dependent plasticity in neuropil volume

Although wild *H. erato* do not have significantly larger absolute volumes for any measured neuropil (Table 4A), differences in allometric scaling or grade-shifts between wild and old insectary-reared individuals are nevertheless evident. Altered scaling affects the MB-ca, Lop, and PB, all of which show shallower scaling relationships (smaller *β*) with rMid in wild-caught individuals (Table 4A; Figure 7B,C). The MB-lo+ped shows an unambiguous grade-shift towards larger size in wild whilst maintaining a common slope, and also shows a major axis shift (Δ_FA_ = 0.250; Fig. 8B1).

**Table 4:**
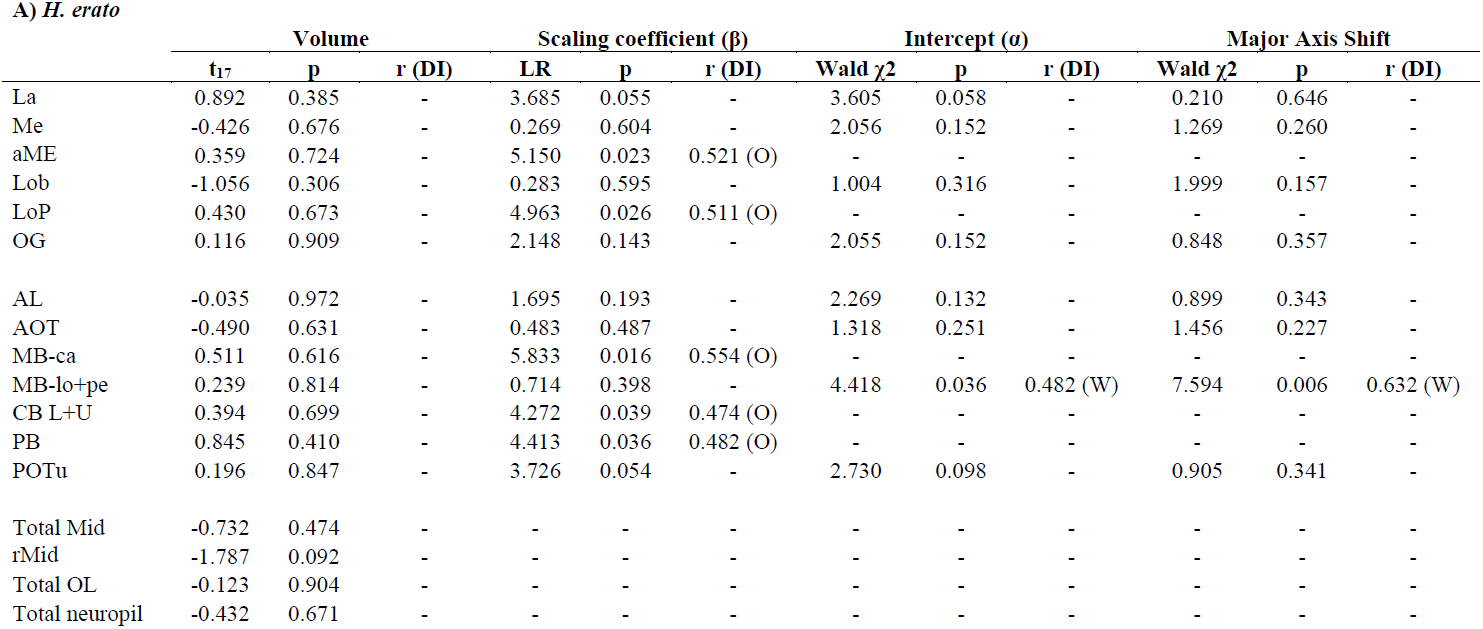
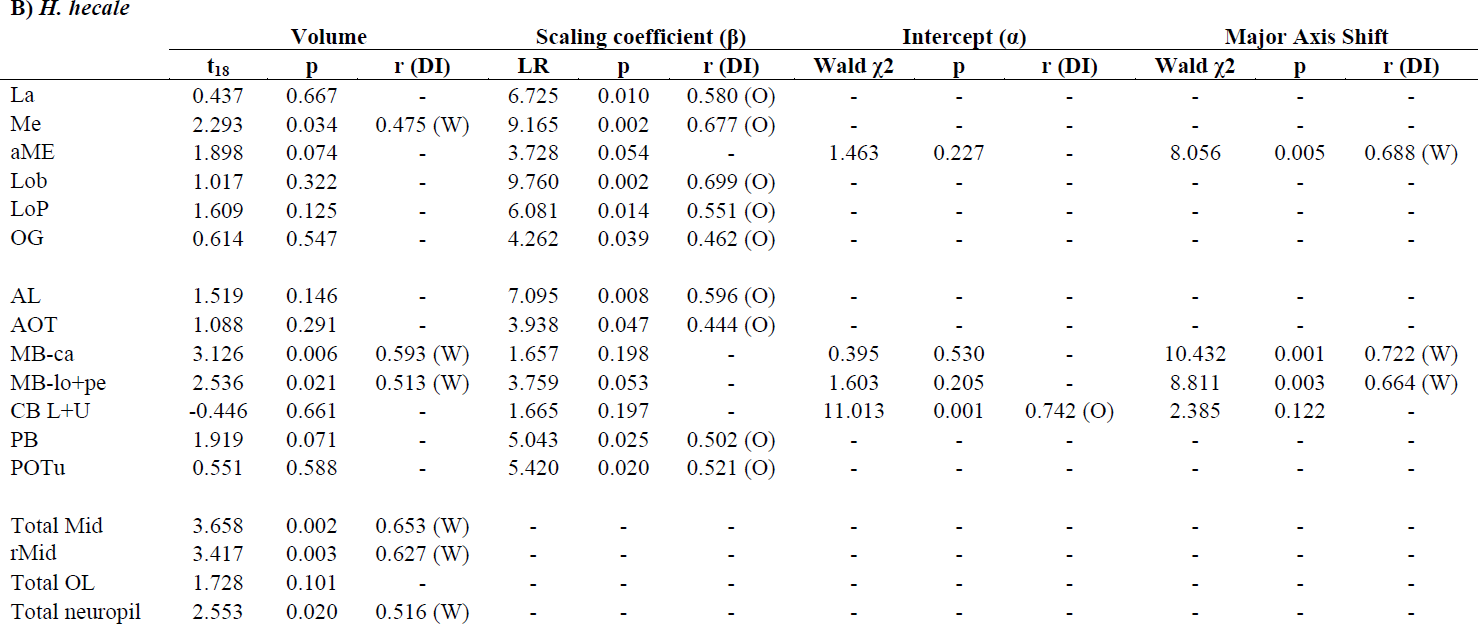
Comparisons between wild caught (W) and old insectary-reared individuals for A) *H. erato* and B) *H. hecale. r* is the effect size. DI indicates the group with a higher value of α, β or fitted axis mean.

In *H. hecale* wild individuals have significantly larger total midbrains (t_18_ = 3.658, p = 0.002). The only segmented neuropil to reflect this difference, however, are the MB-ca and MB-lo+ped (Table 4B; Fig. 8A2,C2), while the rMid is also larger in wild individuals (t_18_ = 3.417, p = 0.003). The average MB-ca volume of old insectary-reared individuals is only 68.3% of the average wild MB-ca volume, for the young insectary-reared individuals it is 49.3% (Figure 8A2,C2). For MB-lo+pe these figures are 76.9% and 58.7% respectively (Figure 8A2,B2). For comparison, in *H. erato* the average MB-ca volume of old insectary-reared individuals is 96.2% of the average wild MB-ca volume, for the young insectary-reared individuals it is 59.7% (Fig. 8A1–C1). For MB-lo+pe these figures are 96.9% and 63.9% respectively (Fig. 8A1–C1).

The only neuropil in the optic lobes to differ significantly in volume in *H. hecale* is the Me. The allometric relationship between neuropil volumes and rMid differs for all neuropil either in the allometric scaling coefficient or the intercept, except for the mushroom body components and aMe (Table 4A; Figure 7E,F). However, for aME this pattern is caused by a lack of allometric scaling in insectary-reared individuals (SMA *p* = 0.552). The mushroom bodies show evidence of a major axis shift along a common slope (MB-ca, Δ_FA_ = 0.355; MB-lo+ped, Δ_FA_ = 0.299; Fig 8B2, C2). Given all grade-shifts result in smaller neuropil volumes relative to rMid volume (Fig. 7E,F) we interpret this as indicating the rMid and mushroom bodies show coordinated environment-dependent increases in volume whilst other neuropil volumes remain largely constant, but with subsequently altered allometric relationships with rMid.

### Allometric scaling of mushroom body components

We further explored the allometric scaling relationships between the three main mushroom body components, the MB-lo and MB-pe (analyzed separately), and the MB-ca. Within wild caught individuals, pairwise comparisons between these structures do not reveal any significant deviation from isometric scaling relationships (test β ≠ 1, p > 0.05). However, the ontogenetic growth we observe between the young and old groups of both species occur through concerted expansion of the MBlo and MBca (i.e. a major axis shift), both of which show grade-shifts in their allometric scaling with the MBpe between the young and old groups (Table 5A). A similar pattern is found comparing *H. hecale* wild and old groups, but there are no significant differences between wild and old *H. erato* with the exception of a narrowly significant difference in the scaling coefficient suggesting MB-lo becomes disproportionally larger as MB-ca increases in wild individuals compared to insectary reared individuals (Table 5B).

**Table 5:** Effects of age (A) and environmental experience (B) on scaling relationships between mushroom body components

| | components | Scaling coefficient ($\beta$) | | | Intercept ($\alpha$) | | | Major Axis Shift | | |
| --- | --- | --- | --- | --- | --- | --- | --- | --- | --- | --- |
| | | Likelihood Ratio | p | r | Wald $\chi^2$ | p | r | Wald $\chi^2$ | p | r (DI) |
| *H. erato* | MB ca + lo | 0.627 | 0.428 | - | 2.249 | 0.134 |  | 16.987 | 0.000 | 0.946 (O) |
|  | MB ca + pe | 1.224 | 0.269 | - | 12.457 | 0.000 | 0.810 | - | - | - |
|  | MB lo + pe | 0.206 | 0.650 | - | 29.286 | 0.000 | 1.000 | - | - | - |
| *H. hecale* | MB ca + lo | 0.100 | 0.752 | - | 0.058 | 0.810 |  | 8.771 | 0.003 | 0.662 (O) |
|  | MB ca + pe | 0.376 | 0.540 | - | 6.422 | 0.011 | 0.567 | - | - | - |
|  | MB lo + pe | 0.118 | 0.731 | - | 5.462 | 0.019 | 0.523 | - | - | - |

B) *Wild vs. old insectary reared*
| | components | Scaling coefficient ($\beta$) | | | Intercept ($\alpha$) | | | Major Axis Shift | | |
| --- | --- | --- | --- | --- | --- | --- | --- | --- | --- | --- |
| | | Likelihood Ratio | p | r | Wald $\chi^2$ | p | r | Wald $\chi^2$ | p | r (DI) |
| *H. erato* | MB ca + lo | 4.083 | 0.043 | 0.464 | 0.139 | 0.709 | - | 0.186 | 0.667 | - |
|  | MB ca + pe | 0.311 | 0.577 | - | 0.732 | 0.392 | - | 0.044 | 0.834 | - |
|  | MB lo + pe | 1.296 | 0.255 | - | 0.213 | 0.645 | - | 0.011 | 0.916 | - |
| *H. hecale* | MB ca + lo | 0.307 | 0.580 | - | 0.398 | 0.528 | - | 7.901 | 0.005 | 0.629 (W) |
|  | MB ca + pe | 2.942 | 0.086 | - | 7.340 | 0.007 | 0.606 | - | - | - |
|  | MB lo + pe | 1.553 | 0.213 | - | 4.086 | 0.043 | 0.452 | - | - | - |

## DISCUSSION

We have described the layout and volume of the major brain neuropils in two species of *Heliconius* butterflies. Our analyses illustrate the role ecology plays in shaping brain structure, and confirm the substantial evolutionary expansion of the *Heliconius* mushroom body first noted by Sivinski (1989). Indeed, our data suggest this previous work underestimated their size. We have further identified neuropil-specific patterns of volumetric variation across young and old insectary-reared and wild individuals that indicate significant post-eclosion growth and experience-dependent plasticity. In the mushroom body, the timing and extent of this ontogenetic plasticity is comparable to that found in insects that strongly rely on spatial memory for foraging (e.g. Withers et al., 1993, 2008; Gronenberg et al., 1996; Fahrbach et al., 1998, 2003; Maleszka et al., 2009; Jones et al., 2013).

### *Interspecific divergence and mushroom body expansion in* Heliconius

Our interspecific analyses across Lepidoptera reveal an unambiguously mosaic pattern of brain evolution (Barton and Harvey, 2000), where the size of individual neuropils deviate from the allometric expectation. Mosaic patterns in mammals, fishes and ants have been interpreted as strong evidence for evolutionary responses to a species’ particular ecological needs (Barton et al., 1995; Huber et al., 1997; Gronenberg and Hölldobler, 1999). In Lepidoptera, this is particularly noticeable in the sensory neuropils (Fig. 6B). The relative volume of the visual neuropils closely reflects diel activity patterns, and the size of the antennal lobes also appears to be strongly associated with a nocturnal or low-light diurnal niche. This is illustrated in a PCA of midbrain neuropil (Fig. 6C) that clusters the olfactorily driven butterfly *G. zavaleta* with night-flying moths (Montgomery and Ott, 2015). Our data further indicate that much of the divergence in AL size among Lepidoptera reflects changes in CFN volume rather than total glomerular volume, implying that changes in the number or branching complexity of AL interneurons dominate over numerical differences in afferent sensory neuron supply, and associated sensitivity. Similarly, the relative constancy in AL glomeruli number indicates that the dimensionality of the afferent coding space is comparable across Lepidoptera with divergent diel patterns (Boeckh and Boeckh, 1979; Rospars, 1983; Berg et al., 2002; Huetteroth and Schachtner, 2005; Masante-Roca et al., 2005; Skiri et al., 2005; Kazawa et al., 2009; Heinze and Reppert, 2012; Carlsson et al., 2013; Montgomery and Ott, 2015).

In contrast with other species-differences that are dominated by changes in the sensory neuropils, *Heliconius* are clearly set apart in our multivariate analysis along an axis heavily loaded by the mushroom bodies. As a percentage of total brain volume, or indeed as a raw volume, *Heliconius* have the largest mushroom body so far reported in Lepidoptera (Sivinski, 1989; Sjöholm et al., 2005; Rø et al., 2007; Kvello et al., 2009; Snell-Rood et al., 2009; Dreyer et al., 2010; Heinze and Reppert, 2012b; Montgomery and Ott, 2015) and one of the largest across insects. This phylogenetic expansion must reflect adaptive change in mushroom body function in response to ecological selection pressures. The derived pollen-feeding behavior of *Heliconius* provides a likely source of this selection (Sivinski, 1989). Several studies have reported this behavior to entail spatially and temporally faithful foraging patterns, guided by visual landmarks (Ehrlich and Gilbert, 1973; Gilbert, 1975, 1993; Mallet, 1986) comparable with the landmark-based trap-lining behavior of foraging bees. Experimental interventions (Mizunami et al., 1998) and comparative neuro-ecological studies (Farris and Schulmeister, 2011) implicate mushroom bodies in visually based spatial memory.

Comparisons across *Heliconius* and non-pollen feeding Heliconiini may provide a test of this spatial memory hypothesis. Sivinski (1989) reported that two individuals of *Dione juno* and *Dryas iulia*, both non-pollen feeding allies to *Heliconius*, had mushroom bodies within the size range of other Lepidoptera. This provides preliminary support that mushroom body expansion coincided with a single origin of pollen feeding at the base of *Heliconius*. However, sampling in a wider range of genera, including the specious *Eueides* which is most closely related to *Heliconius* (Beltrán et al., 2007; Kozak et al., 2015), is required to confirm this conclusion.

Alternative selection pressures also need to be considered, including the degree of host-plant specialization (Brown, 1981) and the evolution of social roosting (Benson, 1972; Mallet, 1986). These factors may well be inter-related, as visits to *Passiflora* may be incorporated into trap-lines between pollen plants (Gilbert, 1975, 1993), and the sedentary home-range behavior required for trap-lining may predispose *Heliconius* to sociality (Mallet, 1986). The latter scenario would parallel the hypothesized origin of sociality in hymenoptera and primates in exaptations of an expanded brain that may have first evolved to support specialization in foraging behavior (Barton, 1998; Farris and Schulmeister, 2011). Regardless of whether pollen feeding provided the initial selection pressure for mushroom body expansion, it is likely that it contributes to meeting the energetic costs of increased neural investment.

### Age- and experience-dependent growth in neuropil volume

In both *H. erato and H. hecale*, the mushroom bodies are significantly larger in aged individuals. Volume increases of 38.0% for the calyx and 34.0% for the lobe system in *H. erato,* and 27.9% for the calyx and 23.7% for the lobes in *H. hecale* are comparable to, if not greater than, the ontogenetic changes seen in Hymenoptera (e.g. c. 30% in *Camponotus floridanus* (Gronenberg et al., 1996); c. 20% in *Bombus impatiens* (Jones et al., 2013)). Our comparisons between aged insectary-reared and wild caught individuals also identify experience-dependent plasticity. This ‘experience’ in the wild likely includes greater range of movement, greater challenges in foraging, and more variable environmental conditions and social interactions.

Our data suggest experience-dependent plasticity particularly affects mushroom body maturation, though the pattern differs between species. In *H. hecale* a strong volumetric difference is found between old insectary-reared and wild caught individuals for both the calyx (32%) and lobes (24%). A concomitant expansion of the unsegmented midbrain results in a pronounced major-axis shift. This is not simply the result of an increased total brain size, however: no other neuropil region shows a comparable increase in wild caught individuals, resulting in widespread grade-shifts in these other neuropils towards smaller size relative to the unsegmented midbrain. This may reflect a coordinated growth between the mushroom bodies and unsegmented midbrain areas or, alternatively, coincident independent expansions. In *erato* old insectary-reared and wild-caught individuals have mushroom bodies of similar absolute size, but allometric grade-shifts over the unsegmented midbrain result in greater relative volumes in wild compared to insectary-reared individuals. The cause of this species difference is unclear, but warrants further investigation.

Finally, it is also notable that plasticity, in particularly age-related growth, is not restricted to the mushroom bodies. Several visual and olfactory neuropils show age- and experience-dependent expansions in *Heliconius,* as they do in other insects (Kühn-Bühlmann and Wehner, 2006; Snell-Rood et al., 2009; Ott and Rogers, 2010; Smith et al., 2010; Heinze and Florman, 2013; Jones et al., 2013). We also find evidence of plasticity in components of the central complex. In *D. plexippus*, size plasticity in the central body and protocerebral bridge has been linked to migratory experience and, by association, the sky compass navigation that supports it (Heinze et al., 2013). The occurrence of similar plasticity in non-migratory butterfly species implies that it may be associated with foraging or locomotor experience more generally, even at much smaller spatial scales.

### Functional relevance of phylogenetic mushroom body expansion

Phylogenetic trends towards larger mushroom bodies involve increases in Kenyon Cell (KC) numbers, clustered into larger numbers of functional sub-units (Farris, 2008). Farris and Roberts (2005) suggest that increasing KC number may provide greater computational capacity by facilitating the processing of more complex combinatorial inputs from afferent projection neurons (Sivan and Kopell, 2004), or through integration across increasingly specialized sub-units (Strausfeld, 2002).

Novel pathways between such specialized KC sub-populations may play an important role in the origin of derived behaviors that require the integration of different sensory modalities (Chittka and Niven, 2009; Strausfeld et al., 2009). Examples of this are provided by Hymenoptera and phytophagous scarab beetles where, in addition to olfactory inputs, the mushroom body calyx receives direct input from the optic lobes (Gronenberg, 2001; Farris and Roberts, 2005; Farris and Schulmeister, 2011). This additional input is reflected in the subdivision of the calyx into the lip, which processes olfactory information, and the collar and basal ring, which process visual information (Gronenberg and Hölldobler, 1999). Visual input to the mushroom bodies has also been demonstrated in some butterflies (Snell-Rood et al., 2009) and moths (Sjöholm et al., 2005) but it has yet to be investigated in *Heliconius*. The *Heliconius* calyx lacks the clear zonation observed in *D. plexippus* that has been suggested to be analogous to the *A. mellifera* lip, collar and basal ring (Heinze and Reppert, 2012). We do not interpret the lack of distinct zonation in *Heliconius* as evidence against functional sub-division, as *Spodoptera littoralis* displays localization of visual processing in the calyx that is not apparent without labeling individual neurons. Given the implied role for visual landmark learning in *Heliconius* foraging behavior (Jones, 1930; Gilbert, 1972, 1975; Mallet, 1986), we hypothesise that their massively expanded mushroom body supports an integration of visual information.

In other species the mushroom body also receives gustatory and mechanosensory input (Schildberger, 1983; Homberg, 1984; Li and Strausfeld, 1999; Farris, 2008). These may also be of relevance in *Heliconius* given the importance of gustatory and mechanosensory reception in host-plant identification (Schoonhoven, 1968; Renwick and Chew, 1994; Briscoe et al., 2013) and pollen loading (Krenn and Penz, 1998; Penz and Krenn, 2000), although it should be noted that there is currently no evidence these behaviors are learnt (Kerpel and Moreira, 2005; Salcedo, 2011; Silva et al., 2014).

### Potential cellular changes associated with ontogenetic mushroom body expansion

The cellular basis of ontogenetic and environmentally induced plasticity may provide further clues as to the functional changes associated with mushroom body expansion during *Heliconius* evolution. The volumetric changes we observe must reflect differences in cell numbers and/or branching and connectivity. It is unknown whether KC neurogenesis is restricted to the larval and pupal stages in Lepidoptera, as it is in Hymenoptera (Fahrbach et al., 1995) where post-eclosion expansion results solely from increased neurite branching (Gronenberg et al., 1996; Farris et al., 2001). In Hymenoptera, age-dependent expansion of the MB-ca accompanies growth of extrinsic neuron processes, whilst increased branching complexity of KCs is associated experience-dependent expansion and foraging specialization in social castes (Farris et al., 2001; Jones et al., 2009). This suggests that changes in the calyx circuitry involving increased synaptic connections onto KC dendrites may be responsible for the volumetric changes associated with behavioral experience. Although we have not yet measured KC number or dendritic branch length, the grade-shifts of MB-ca and MB-lo over MB-pe, that are uncovered by our allometric analyses, indicate that post-eclosion growth is not solely (if at all) due to additional KCs. This is because each additional KC will necessarily contribute volume to all three major MB compartments, MB-ca, MB-pe and MB-lo. Indeed, Ott and Rogers (2010) proposed that wiring overheads might increase non-linearly with increasing KC numbers to explain the positive allometric scaling of MB-ca over MB-lo observed in some other insects. The resultant scaling need not be isometric, however, this effect will always produce a constant allometric scaling relationship. In contrast, the grade-shifts we observe are most likely explained by increased dendritic growth and connectivity in the MB-ca and MB-lo. Confirming this interpretation, and understanding its functional relevance, may provide some insight into how environmental information is stored during post-eclosion development.

## Conclusions

Our volumetric analyses uncover extensive phylogenetic expansion and ontogenetic plasticity of *Heliconius* mushroom bodies. Both processes may be linked to the derived foraging behavior or this genus, which relies on allocentric memory of pollen resources (Gilbert, 1975; Sivinski, 1989). This hypothesis must now be confirmed in wider comparative analyses and tested explicitly in behavioral experiments. Our phenotypic observations furthermore provide the necessary framework for analyses of the underpinning neuronal mechanisms regulating neuropil size, and of the consequences for circuit function.

## Acknowledgments

The authors are grateful to Adriana Tapia, Moises Abanto, William Wcislo, Owen McMillan, Chris Jiggins, and the Smithsonian Tropical Research Institute for assistance, advice, and the use of the *Heliconius* insectaries at Gamboa, Panama, and the Ministerio del Ambiente for permission to collect butterflies in Panama. We also thank Judith Mank’s research group at UCL for helpful advice and feedback, and the UCL Imaging Facility for help with confocal microscopy.

## Conflict of interest statement

The authors declare no conflict of interest.

## Role of authors

All authors read and approved the final manuscript. Study conception: SHM. Study design and preliminary experiments: SHM, RMM, SRO. Fieldwork and insectary rearing: SHM, RMM. Acquisition of data, analysis, interpretation and initial manuscript draft: SHM. Final interpretation and drafting: SHM, RMM, SRO.

## Financial support

This research was supported by research fellowships from the Royal Commission of the Exhibition of 1851 and Leverhulme Trust, a Royal Society Research Grant (RG110466) and a British Ecological Society Early Career Project Grant awarded to SHM. RMM was supported by a Junior Research Fellowship from King’s College, Cambridge and an Ernyst Mayr Fellowship from STRI. SRO was supported by a University Research Fellowship from the Royal Society, London (UK).

## Abbreviations

AL: antennal lobe
aMe: accessory medulla
AN: antennal nerve
AOTu: anterior optic tubercule
CB: central body
CBL: lower central body
CBU: upper central body
CFN: central fibrous neuropil of AL
DMSO: dimethyl suphoxide
Glom: glomeruli
HBS: HEPES-buffered saline
iMe: inner medulla
iRim: inner rim of the lamina
KC: kenyon cell
La: lamina
LAL: lateral accessory lobes
Lo: lobula
LoP: lobula plate
LU: lower unit of AOTu
MB: mushroom body
MB-ca: mushroom body calyx
MB-lo: mushroom body lobes
MB-pe: mushroom body peduncle
MB-lo+pe: mushroom body lobes and peduncle combined
MBr: midbrain
Me: medulla
MGC: macro-glomeruli complex
NGS: normal goat serum
no: noduli
NU: nodule unit of AOTu
oMe: outer medulla
OR: olfactory receptor
OGC: optic glomerular complex
PA: pyrrolizidine alkaloids
PB: protocerebral bridge
PC: principal component
POTu: posterior optic tubercle
rMid: rest of midbrain
SP: strap of AOTu
UU: upper unit of AOTu
ZnFA: Zinc-Formaldehyde solution

